# Rear cortex contraction aids in nuclear transit during confined migration by increasing pressure in the cell posterior

**DOI:** 10.1101/2022.09.10.507419

**Authors:** Jeremy Keys, Brian C.H. Cheung, Margaret A. Elpers, Mingming Wu, Jan Lammerding

**Author notes:** Correspondence should be addressed to Jan Lammerding.

## Abstract

As cells migrate through biological tissues, they must frequently squeeze through micron-sized constrictions in the form of interstitial pores between extracellular matrix fibers and/or other cells. Although it is now well recognized that such confined migration is limited by the nucleus, which is the largest and stiffest organelle, it remains incompletely understood how cells apply sufficient force to move their nucleus through small constrictions. Here, we report a mechanism by which contraction of the cell rear cortex pushes the nucleus forward to mediate nuclear transit through constrictions. Laser ablation of the rear cortex reveals that pushing forces behind the nucleus are the result of increased intracellular pressure in the rear compartment of the cell. The pushing forces behind the nucleus depend on accumulation of actomyosin in the rear cortex and require Rho-kinase (ROCK) activity. Collectively, our results suggest a mechanism by which cells generate elevated, intracellular pressure in the posterior compartment to facilitate nuclear transit through 3D constrictions. This mechanism may supplement or even substitute for other mechanisms supporting nuclear transit, ensuring robust cell migrations in confined 3D environments.

## Introduction

The migration of cells through tissues is essential to many biological processes, including embryogenesis, immune surveillance, wound healing, and cancer metastasis (Doyle et al., 2013; Yamada and Sixt, 2019). During in vivo migration, cells must navigate through extracellular matrix networks and layers of endothelial cells, which form constrictions as small as 1 to 2 µm in diameter (Bone et al., 2016; Khatau et al., 2012; Weigelin et al., 2012; Wolf et al., 2013), i.e., substantially smaller than the size of the cell and the nucleus. Although cells can readily alter their morphology through cytoskeletal reorganization to adjust to the size of the constriction, the deformation of the nucleus represents a unique physical challenge, as it is the largest and stiffest organelle in the cell (Caille et al., 2002; Denais et al., 2016; Khatau et al., 2012; McGregor et al., 2016; Wolf et al., 2013). Because of the size and stiffness of the nucleus, cells must generate substantial intracellular forces to deform the nucleus through the constrictions. The process of deforming and moving the nucleus through these narrow spaces, henceforth referred to as “nuclear transit,” limits the rate at which cells migrate in confined 3D environments (Davidson et al., 2015; Denais et al., 2016; Harada et al., 2014; Jayo et al., 2016; McGregor et al., 2016; Rowat et al., 2013; Thomas et al., 2015; Wolf et al., 2013; Yadav et al., 2018).

Various mechanisms have been proposed to explain how cells accomplish nuclear transit through confined environments (Marks and Petrie, 2022; McGregor et al., 2016). In neurons and developing muscle cells, microtubule associated motors kinesin and/or dynein are crucial for nuclear movement by applying forces to the nucleus via the LINC complex (Kalukula et al., 2022; Tsai et al., 2010; Wilson and Holzbaur, 2012; Wu et al., 2011; Zhu et al., 2017). In other cell types, nuclear movement is primarily driven by actomyosin mediated forces. One commonly described mechanism suggests that actomyosin and intermediate filaments at the front of the cell pull the nucleus forward (Davidson et al., 2020; Jayo et al., 2016; Petrie et al., 2014). Other studies have implicated pushing forces in the process, mediated through perinuclear actomyosin fibers, which wrap around the nucleus (Thomas et al., 2015), or Arp2/3-mediated perinuclear actin polymerization (Thiam et al., 2016). Each of these mechanisms depend on physical connections between the nucleus and cytoskeleton. Another proposed mechanism suggests that the nucleus is pushed forward by contraction of actomyosin in the rear cortex of migrating cells (Hetmanski et al., 2019; Ju et al., 2023; Lämmermann et al., 2008; Lee et al., 2021; Mistriotis et al., 2019; Poincloux et al., 2011). Support for the nucleus being pushed from behind derives from the observation that cells migrating in 3D environments commonly accumulate actin in the trailing edge of the cell, and that contraction of the cell rear often precedes nuclear transit (Hetmanski et al., 2019; Lämmermann et al., 2008; Mistriotis et al., 2019; Thiam et al., 2016). However, functional evidence for this mechanism is still lacking, including how forces from the contraction of the rear cortex are transmitted to the nucleus. Lastly, many cell lines have shown the ability to migrate using multiple modes, such as adhesion-independent “amoeboid” migration and integrin adhesion-based “mesenchymal” migration, depending on cellular contractility and environmental adhesiveness (Liu et al., 2015; Sahai and Marshall, 2003). This plasticity of migration modes suggests that it is possible that cells may similarly switch between nuclear transit mechanisms in differing biological contexts (Marks and Petrie, 2022), but experimental evidence for this has been lacking. Thus, despite extensive previous research, several questions remain regarding the potential mechanism cells use to apply cytoplasmic forces to the nucleus during confined migration, including (1) how forces are transmitted from the contracting cell rear to the nucleus, and (2) to what extent contraction of the cell rear contributes to or is necessary for nuclear transit through small constrictions. Using microfluidic devices that mimic confined interstitial spaces, along with advanced imaging and cellular manipulation approaches, we here describe a mechanism by which actomyosin in the rear cortex contracts to push the nucleus forward through constrictions. This mechanism is associated with enrichment of actin and active myosin in the rear cell cortex and depends on elevated RhoA-ROCK signaling. RhoA-ROCK mediated contraction of the rear cortex results in elevated intracellular pressure in the posterior compartment of the cell, which pushes the nucleus from behind to support migration through confined spaces.

## Results

### Contraction of the posterior actin cortex promotes nuclear transit through constrictions

To model cell migration through confined environments, we seeded cells into microfluidic devices that enable the close observation of nuclear transit through precisely defined constrictions at high spatiotemporal resolution (Davidson et al., 2015; Keys et al., 2018). In these devices, cells migrate through collagen-coated channels with constrictions measuring 1 to 2 µm in width and 5 µm in height (i.e., cross-sectional area ≤2×5 µm^2^), formed by poly-dimethyl siloxane (PDMS) pillars (**Fig. 1A**). The devices also feature wider control channels (cross-sectional area 15×5 µm^2^) that do not require substantial nuclear deformation during migration (**Fig. 1A**) (Davidson et al., 2015). The size of the narrow constrictions reflects the geometries of small pore sizes that migrating cancer cells encounter in vivo (Weigelin et al., 2012), while the width of the control channels was chosen to be wider than the typical diameter of nuclei (Lammerding, 2011) and corresponds to the upper range of interstitial spaces (Weigelin et al., 2012). We have previously demonstrated that the 5 µm height is sufficient to vertically confine migrating cells, and that cells adhere to both top and bottom surfaces of the cell, plus the vertical walls (Davidson et al., 2015, 2014), thereby creating true 3D confined environments that model confined migration *in vivo* (Denais et al., 2016; Wang et al., 2021).These devices provide several advantages over imaging cells migrating through collagen matrices, as they enable long-term imaging of cells migrating along a single focal plane, through precisely defined constriction geometries. In contrast, in vitro assembled collagen matrices have heterogeneous, randomly distributed constriction geometries and often lead to cells migrating out of the imaging plane during prolonged image acquisition series.

**Figure 1.**
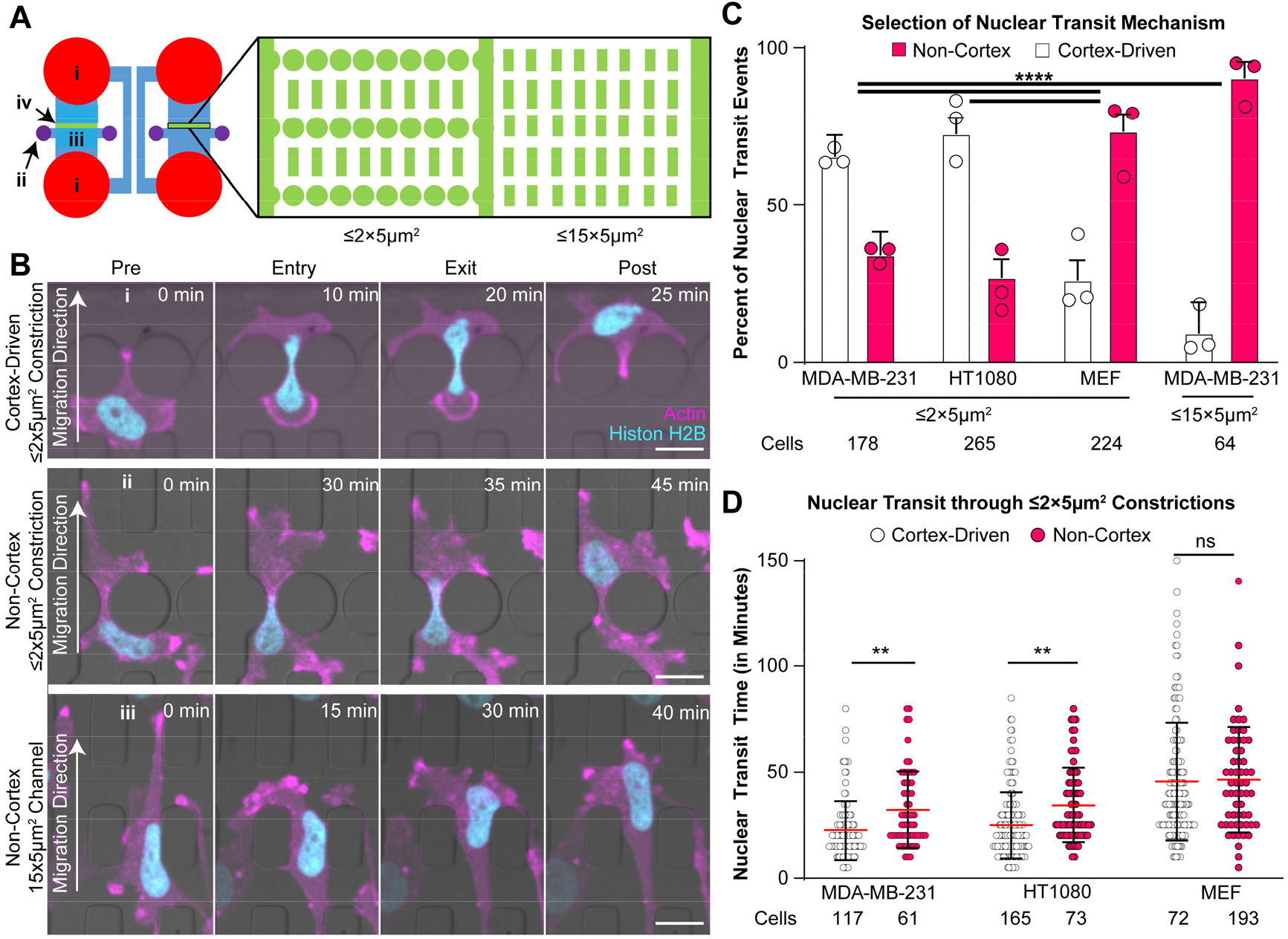
Cortex-driven nuclear transit provides an advantage to cells migrating through narrow constrictions. (**A**) Schematic layout of microfluidic device used to study confined migration through precisely defined constrictions. (i) Reservoirs for cell media; (ii) seeding ports to add cells; (iii) seeding area where cells attach prior to migration through constrictions; (iv) constriction area, shown in detail in inset, to the right. The inset shows areas with constrictions (left) and the larger control channels (right), with the corresponding cross-sections listed below each area. (**B**) Representative image sequences of MDA-MB-231 cells co-expressing fluorescently labeled histones (mNeonGreen-H2B, cyan) and actin (mCherry2-actin chromobody, magenta). MDA-MB-231 cells adopt varying morphological characteristics as the nucleus squeezes through microfluidic constrictions using either (i) a rear cortex-driven mechanism, or (ii) a non-cortex driven mechanism. Scale bar: 20 µm. (**C**) Relative frequency of cortex-driven and nuclear transit mechanisms in varying cell types migrating through either narrow constrictions (≤2×5 µm^2^) or wider control channels (15×5 µm^2^). Error bars represent 95% confidence intervals. ****, *p < 0*.*0001*, based on a Chi-Squared test of independence. Circles represent means of experimental replicates. (**D**) Nuclear transit times through ≤2×5 µm^2^ constrictions of different cell types, subdivided into cortex-driven and non-cortex driven nuclear transit mechanisms. Red line represents sample mean. Error bars represent standard deviation. **, *p* < 0.01; ****, *p* < 0.0001, based on one-way ANOVA.

We chose to study MDA-MB-231 metastatic breast cancer cells because they exhibit the ability to switch between different migration modes (Geiger et al., 2019; Sahai and Marshall, 2003) and therefore may use multiple mechanisms to drive nuclear transit. MDA-MB-231 cells stably expressing fluorescently labeled histones (mNeonGreen-histone H2B (Davidson et al., 2015)) and actin (mCherry2-actin chromobody (Chromotek)) exhibited two distinct patterns of actin accumulation around the nucleus while migrating through constrictions in microfluidic migration devices. Most cells formed a rounded, actin-enriched cortex in the rear of the cell, the contraction of which was associated with the nucleus squeezing through the constriction (**Fig. 1B-i, Supp. Movie 1, Supp. Fig. 1B-F**). In other cells, the nucleus transited through the constriction without the cell forming a rounded rear cortex (**Fig. 1B-ii, Supp. Fig. 1G-I**). Notably, whereas the formation of a distinct, rounded rear cortex was observed in the majority of MDA-MB-231 cells migrating through narrow constrictions, it was rarely observed in cells migrating through the larger control channels, nor in cells migrating on 2D glass substrates (**Fig. 1B-iii, 1C, Supp. Fig. S2C**). We observed similar patterns of actin accumulation at the rear of MDA-MB-231 cells migrating through 3-dimensional collagen matrices (**Supp. Fig. S2D-E**). Together with previous observations of the formation of a rounded rear cortex in MDA-MB-231 cells during transendothelial migration (Chen et al., 2016), our results indicate that these nuclear transit mechanisms are relevant during migration in physiological environments.

We hypothesized that the differences in cytoskeletal organization prior to and during nuclear transit indicate two distinct nuclear transit mechanisms, which may correspond to the previously described “pushing” and “pulling” forces. Accumulation of actin in the rear cortex suggests a “cortex-driven” pushing of the nucleus from behind, whereas accumulation of actin in front of the nucleus suggests “non-cortex-driven” pulling at the front of the nucleus. We recorded similar actin dynamics in HT1080 fibrosarcoma cells in microfluidic migration devices (**Fig. 1C, Supp. Fig. S2B**), another cancer cell line that shows considerable plasticity in its migratory mode (Petrie et al., 2016; Wolf et al., 2003). In contrast, mouse embryonic fibroblasts (MEFs), which tend to use a mesenchymal migration mode (Gadea et al., 2007), predominantly used a non-cortex driven nuclear transit mode (**Fig. 1C, Supp. Fig. S2A, Supp. Movie 2**).

Based on these observations, we hypothesized that the rear cortex contraction may aid MDA-MB-231 and HT1080 cells in nuclear transit through narrow constrictions by applying “pushing” forces to the nucleus from the back of the cell. To determine whether the rear cortex contraction produced more rapid nuclear transit, we compared the transit times between cells using either (rear) cortex-driven or non-cortex driven nuclear transit for each of the cell lines. In MDA-MB-231 and HT1080 cells, cortex-driven cells translocated their nuclei significantly faster through narrow constrictions than non-cortex-driven cells (**Fig. 1D**), suggesting that in these cell types, the rear cortex contractions provide an advantage over non-cortex driven migration. In contrast, we did not find any differences in nuclear transit time between MEFs using the cortex versus the non-cortex driven mechanism (**Fig. 1D**). The preference towards a cortex-driven mechanism seen in MDA-MB-231 and HT1080 cells, in contrast to MEFs, may be due to the greater migratory plasticity that is commonly observed in these cancer cell lines (Sahai and Marshall, 2003). As MDA-MB-231 and HT1080 cells are capable of migrating using both mesenchymal and amoeboid migration modes, depending on environmental and biological factors, they may be able to use this additional cortex-driven nuclear transit mechanism to support passage through narrow constrictions. MEFs, in contrast, are more conventionally mesenchymal in their migration (Gadea et al., 2007), and therefore may be less able to use this additional migration mechanism. Taken together, these results suggest that rear cortex contraction may act as an alternative or supplemental mechanism in confined migration, which enables more rapid migration through constrictions but is not necessarily required for nuclear transit.

Since previous studies had suggested that pulling forces at the leading edge of the nucleus require connections between the nucleus and cytoskeleton (Davidson et al., 2020; Petrie et al., 2014), we investigated whether disruption of the linker of nucleoskeleton and cytoskeleton (LINC) complex caused a switch to a cortex-driven nuclear transit mechanism. We therefore chose to use MEFs as a model system, as we had previously observed them to primarily use a non-cortex driven migration mode. MEFs in which endogenous Nesprin-2 was tagged with GFP (GFP-Nesprin-2, Davidson et al. 2020) and expressing an mCherry2-Actin chromobody were modified to additionally express either a blue fluorescent protein (BFP) or a BFP-tagged dominant-negative KASH construct (DN-KASH), which binds with Sun proteins at the nuclear envelope and prevents the association between Sun proteins and endogenous Nesprin proteins, thereby disrupting force transmission between the cytoskeleton and nucleus (Lombardi et al., 2011). Whereas GFP-Nesprin-2 was localized to the nuclear envelope in non-modified cells and in cells expressing a BFP-only control constructs, in DN-KASH expressing MEF cells, Nesprin-2 was mis-localized from the nuclear envelope (**Supp. Fig. S3A-B)**, confirming disruption of the LINC complex by the DN-KASH construct. Surprisingly, LINC complex disruption did not correspond with increased nuclear transit times, nor in a shift towards a cortex-driven mechanism (**Supp. Fig. S3C-D**).

To determine if these trends were cell-line specific, we modified MDA-MB-231 cells to express either the doxycycline inducible DN-KASH construct or a BFP-only control. Similar to the results in the MEFs, DN-KASH expression resulted in endogenous nesprins (here, Nesprin-1) being mislocalized from the nuclear envelope (**Supp. Fig. 3E-F**). DN-KASH expression did not significantly alter the proportion of cells employing cortex-driven vs. non-cortex driven migration modes (**Supp. Fig 3G**), consistent with the results in MEFs (**Supp. Figs. S3D**). However, whereas DN-KASH expression had no effect on the nuclear transit times of MDA-MB-231 cells using cortex-driven migration, DN-KASH expression slowed the nuclear transit rate of MDA-MB-231 cells with non-cortex driven migration (**Supp. Fig. 3H**). This increase in nuclear transit time upon LINC complex disruption differed from the results in MEFs, where LINC complex disruption had no effect on nuclear transit time (**Supp. Fig. 3C**). Collectively, these findings indicate that although some non-cortex driven cells may require the LINC-complex for efficient nuclear transit, the LINC complex is dispensable for cortex driven nuclear transit. As this effect was only observed in one of the two tested cell lines, the precise requirements of the LINC complex for non-cortex driven nuclear transit remains to be explored in future studies.

### Rear cortex driven nuclear transit is associated with actin and myosin II accumulation at the rear cortex and dependent on RhoA-ROCK activity

We hypothesized that cortex-driven cells localize contractile machinery to the rear cortex to push the nucleus through constrictions. Previous studies found that non-muscle myosin II is important for force generation during 3D confined migration (Davidson et al., 2020; Thomas et al., 2015); thus, we performed timelapse video microscopy on MDA-MB-231 cells transfected to transiently express GFP-myosin IIA and mCherry-myosin IIB in the migration devices. Both myosin II isoforms localized to the rear cortex of cells as the nucleus squeezed through constrictions (**Fig. 2A, Supp. Fig. 1J-Q**). Notably, the accumulation of myosin II at the rear cortex was only observed in cells migrating through narrow constrictions, whereas in cells migrating through larger control channels or on 2D surfaces, i.e., in situations not utilizing rear-cortex driven nuclear transit, myosin II was localized to the front of the cell body or the leading edge of the nucleus (**Fig. 2A, Supp. Fig. S1R-S**). Our studies also revealed that individual cells can switch between non-cortex-driven and cortex-driven mechanisms during nuclear transit. In some instances, cells that entered constrictions with GFP-myosin II initially primarily localized at the leading edge of the nucleus failed to pass through constrictions. When the nuclear transit attempt was not successful, the cells backed out of the constriction, and re-entered the constriction with GFP-myosin IIA localized to the rear cortex, using rear cortex contraction to successfully push the nucleus through the constriction (**Supp. Movie 3**). These findings suggest that cells may actively switch between nuclear transit modes while squeezing through small constrictions. In our analysis, out of 40 cells that completed multiple nuclear transits within the recorded timelapse sequence, 9 cells (22.5%) “switched” between nuclear transit mechanisms during successive nuclear transit events.

**Figure 2:**
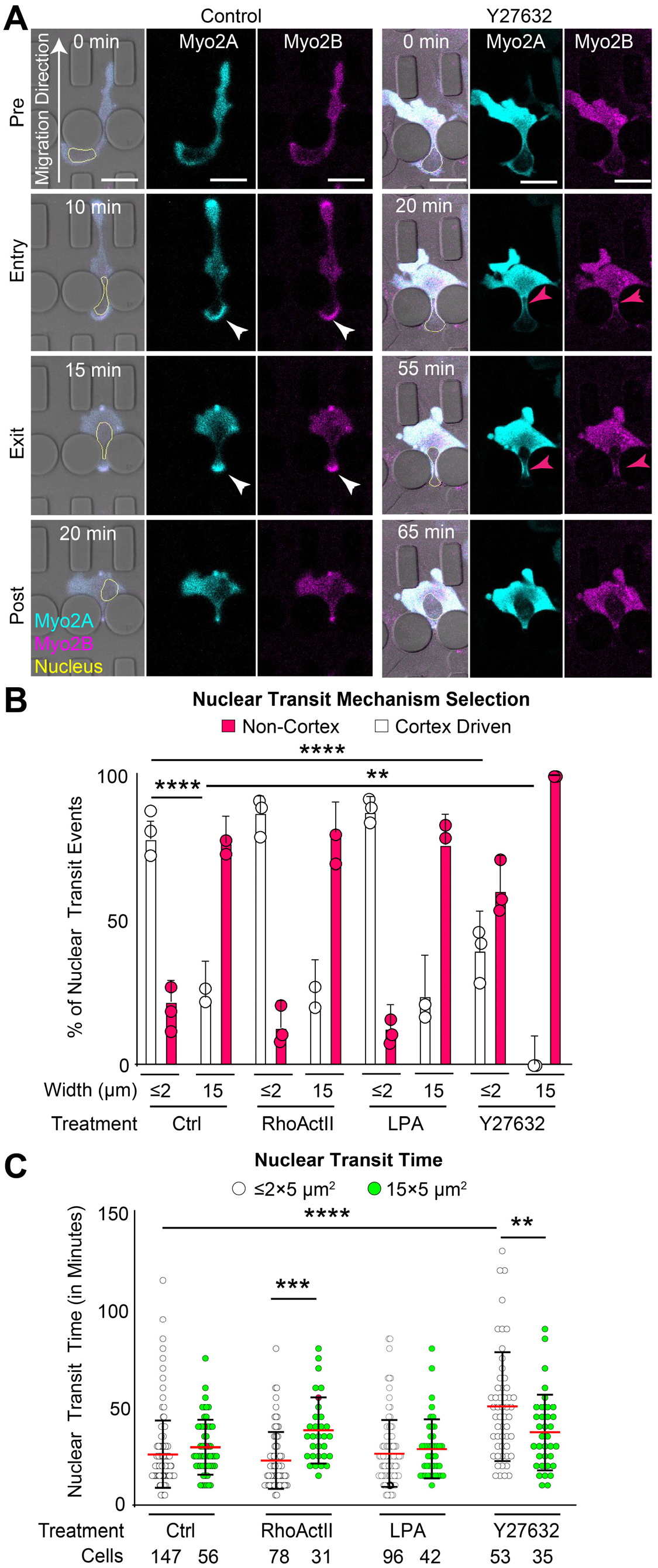
Rear cortex driven nuclear transit is associated with localization of myosin II to the rear cortex. (**A**) Representative time-lapse series of MDA-MB-231 cells expressing GFP-myosin IIA and mCherry-myosin IIB migrating through microfluidic migration devices. White arrows indicate accumulation of myosin II at the rear cortex. Magenta arrows point to accumulation in front of the nucleus. The nucleus is outlined in yellow for better visibility in the merged image. Scale bars: 20 µm (**B**) Percentage of nuclear transit events using cortex-driven or non-cortex driven modes in either ≤2×5 µm^2^ or 15×5 µm^2^ constrictions following treatment with either RhoA activators (RhoActII or LPA), ROCK inhibitor (Y27632), or vehicle control. Error bars represent 95% confidence intervals. **, *p* < 0.01; ****, *p* < 0.0001, based on a Chi-Squared Test of Independence. Circles represent mean of each experimental replicate. (Number of cells analyzed per group: *n* = 147, 56, 78, 31, 106, 42, 53, 35, respectively). (**C**) Nuclear transit times of MDA-MB-231 cells migrating through constrictions following treatment with RhoA activators (RhoActII or LPA) or ROCK inhibitor (Y27632). Red line represents mean. Error bars represent standard deviation. **, *p* < 0.01; ***, *p* < 0.001; ****, *p* < 0.0001, based on a one-way ANOVA. Number of cells in each group indicated below the graphs.

Since myosin contractility is regulated through RhoA and Rho-associated protein kinase (ROCK) (Ridley et al., 2003), we hypothesized that cells locally activate RhoA in the rear cortex to generate actomyosin contractile forces to push the nucleus from behind. To record the spatiotemporal dynamics of RhoA activation during nuclear transit, we stably modified MDA-MB-231 cells with a previously established FRET-based biosensor for RhoA activity (Pertz et al., 2006) and imaged the cells as they migrated through constrictions. Local RhoA activity was determined by measuring the average FRET ratio in either the rear cortex, front cortex, or directly in front of the nucleus (**Supp. Fig. S4A**). Since levels of the RhoA sensor inside the nucleus were very low, as expected, which could negatively affect the robustness of ratiometric FRET analysis, the nucleus was masked-out and excluded from the analysis (**Supp. Fig. S4A**). FRET ratios of the RhoA biosensor were evaluated prior to nuclear transit (“Pre”), during nuclear entry into a constriction (“Entry”), immediately upon nuclear exit from the constriction (“Exit”), and after the cell had left the constriction (“Post”) (**Fig. 3A**). Consistent with prior reports (Pertz et al., 2006), RhoA activity was increased at the leading edge of the migrating cells (**Fig. 3A**). However, as cells migrated through narrow constrictions, RhoA activity in the rear cortex became elevated relative to other regions within the cell during nuclear transit (**Fig. 3B**). This increase in RhoA activity in the rear cortex was unique to cells migrating through narrow 3D constrictions and was not observed in cells migrating through the wider control channels or on 2D substrates (**Supp. Fig. S4B**). In confined conditions, RhoA activity in the rear cortex increased as the nucleus entered the constriction, and then returned to baseline levels during the later stages of nuclear transit (**Fig. 3C**). Unlike RhoA activity at the rear cortex, RhoA activity at the leading edge of the cell remained constant throughout nuclear transit (**Supp. Fig. S4D**). RhoA activity in front of the nucleus (**Supp. Fig. S4A, yellow zone**) did not increase upon nuclear entry into the constriction but increased at the end of nuclear transit (**Supp. Fig. S4C**). Collectively, these observations suggest a sequential process in which RhoA is first activated at the posterior of the cell, causing contraction of actomyosin in the rear cortex to push the nucleus into a constriction, followed by activation of RhoA in front of the nucleus to contract actomyosin at the leading edge of the nucleus to pull the nucleus through the constriction.

**Figure 3.**
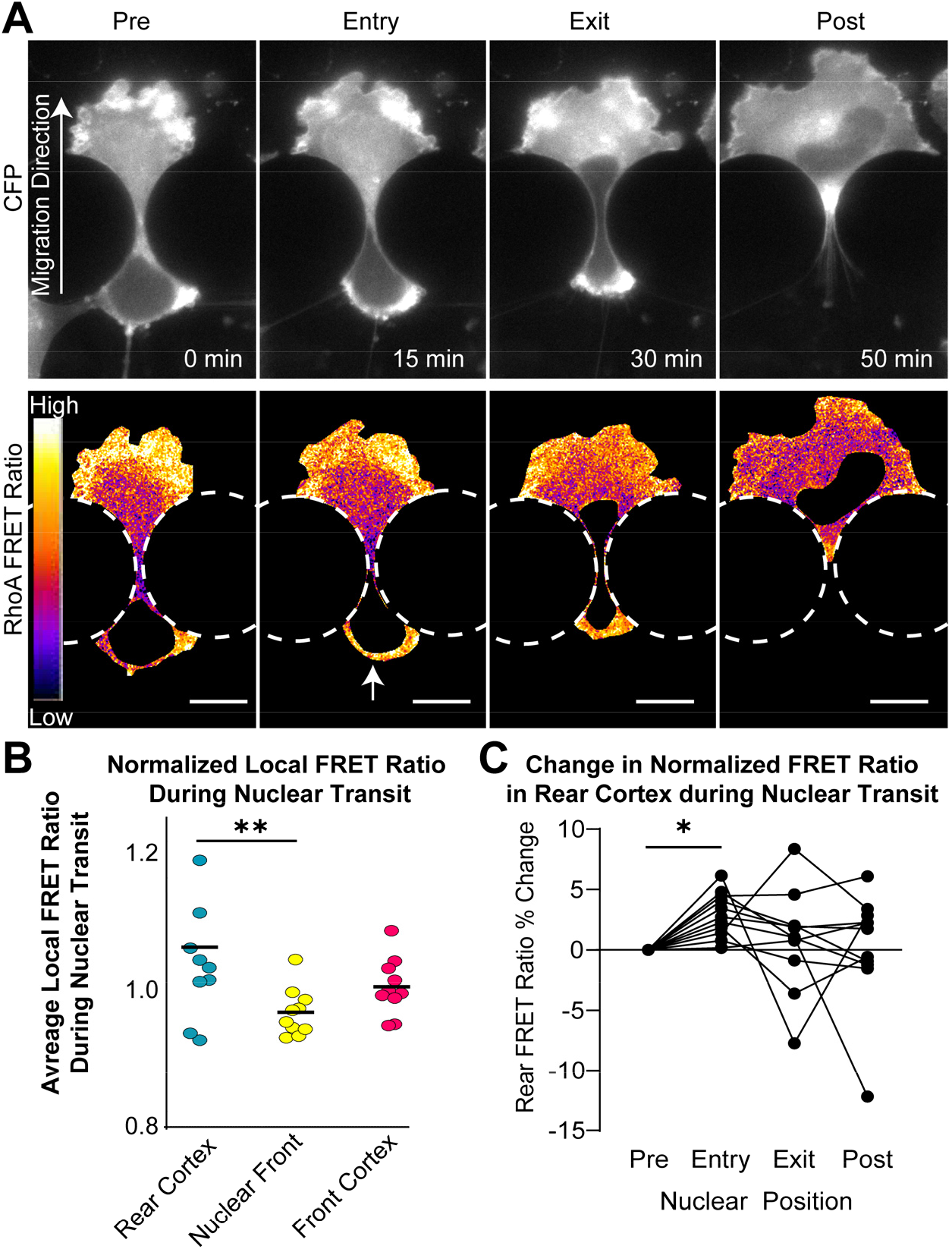
Cortex-driven nuclear transit is associated with increased RhoA activity at the rear cortex. (**A**) Representative time lapse series of FRET ratio of MDA-MB-231 cell expressing RhoA-FRET biosensor migrating through a microfluidic constriction. The bottom series shows the RhoA-FRET ratio during nuclear transit, illustrating increased RhoA activity at the rear cortex (arrow) and the leading edge of the cell. Scale bars: 20 µm (**B**) Comparison of RhoA activity at different intracellular locations based on the recorded RhoA-FRET values for cells migrating through ≤2×5 µm^2^ constrictions. Each data point represents local RhoA FRET measurements averaged across all time points during a single nuclear transit event (*n* = 10 cells/group). Black lines represent the mean for each group. **, *p* < 0.01 using Friedman test. (**C**) Time course of FRET ratio within rear cortex of individual cells (normalized to average FRET ratio of whole cell) at each phase of nuclear transit through ≤2×5µm^2^ constrictions. *, *p* < 0.05 using Friedman test.

As the cortex-driven nuclear transit mechanism was associated with active RhoA and myosin II at the rear cortex, we hypothesized that the cortex-driven nuclear transit mechanism depends on Rho-ROCK activity. We treated cells with either ROCK inhibitor (Y27632) or RhoA activators (LPA or RhoA Activator II) and measured nuclear transit times in the microfluidic devices. ROCK inhibitor treatment significantly slowed nuclear transit through the constrictions but did not slow migration through unconfined channels (**Fig. 2C**), suggesting that ROCK activity is particularly important for confined migration. Interestingly, ROCK inhibition increased the frequency of non-cortex driven nuclear transit events and altered localization of myosin IIA and IIB during nuclear transit, causing myosin to localize towards the front of the nucleus, rather than in the rear cortex as is seen in vehicle-only control cells (**Fig. 2A-B**). As ROCK inhibition slowed down nuclear transit through narrow constrictions but not through unconfined channels, these findings also suggests that the role of ROCK in pushing the nucleus from behind is unique to nuclear transit through narrow constrictions. This observation mirrors our earlier observation that cells in unconfined channels rarely use a cortex-driven migration mode. Ectopic activation of RhoA with either LPA or RhoA Activator II had no effect on the nuclear transit rate or nuclear transit mechanism in MDA-MB-231 cells (**Fig. 2C**), presumably due to the already abundant levels of LPA present in the serum. The lack of effect of pharmacological RhoA activators on nuclear transit rate or mechanism may indicate that RhoA in these cells was already highly activated by serum in the cell media, or that global RhoA activation without spatial control within the cell is insufficient to activate the cortex-driven transit mechanism. Although actomyosin contractility likely plays a role in both cortex-driven and non-cortex driven mechanisms, these results further suggest that cortex-driven nuclear transit requires a particularly high level of Rho-ROCK mediated contractility in contrast to non-cortex driven nuclear transit.

### Laser ablation of the cell cortex reveals a pressure-driven nuclear transit mechanism

Having established that cortex-driven cells localize actomyosin and active RhoA to the rear cortex at the beginning of nuclear transit, we hypothesized that contraction of the rear cortex pressurizes the rear compartment of the cell relative to the front compartment, pushing the nucleus forward through the constriction. To probe differences in intracellular pressure generated by rear cortex contraction, we used a two-photon laser to ablate specific cytoskeletal regions of migrating cells while they squeezed their nucleus through microfluidic constrictions (**Fig. 4A; Supp. Fig. S5A)**. As the nucleus transits through a constriction, the cytoskeletal forces pushing or pulling on the nucleus are counteracted by the resistance of the constriction in the form of the friction and the normal forces at the interface between the nucleus and the constriction (**Supp. Fig. S5B-C**).

**Figure 4.**
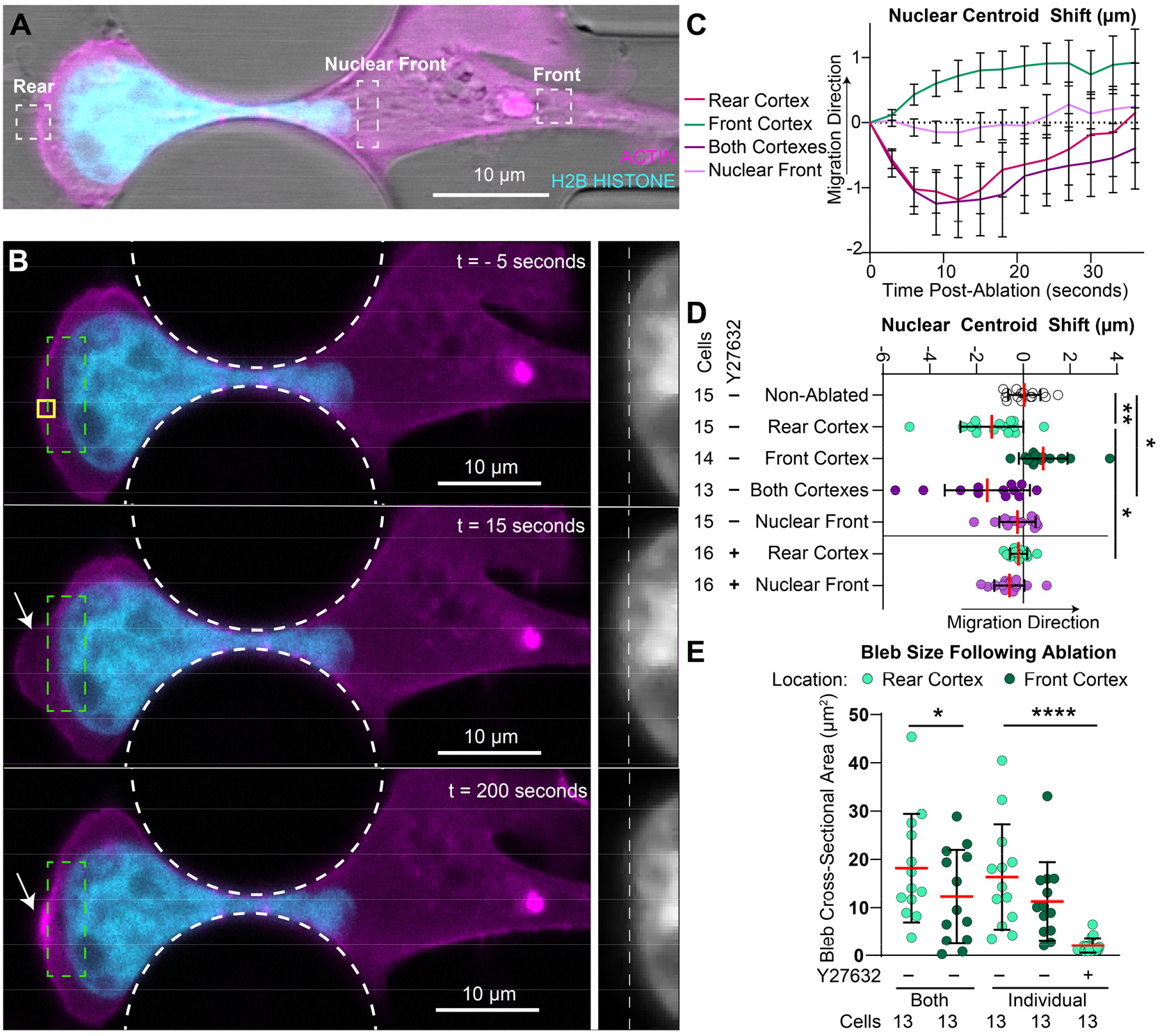
Nuclear movement following laser ablation of local actin cytoskeleton reveals that cortex-driven nuclear transit results from increased pressure in the rear compartment. (**A**) Laser ablation set-up. Dashed boxes represent regions targeted for laser ablation in cells starting to undergo nuclear transit through a small (≤2×5 µm^2^) constriction. Cells were modified to stably express mNeonGreen-histone H2B and mCherry actin-chromobody to visualize nuclear movement and actin cytoskeleton, respectively. (**B**) Representative time lapse series of cell ablated at the rear cortex. Yellow box represents laser ablation target. Arrow shows plasma membrane bleb formation post-ablation. Dashed green box corresponds with inlay of mNeonGreen-histone H2B signal on right. In inlay region: dashed white line represents starting point of leading edge of nucleus. (**C**) Trace of nuclear centroid shift following laser ablation. Error bars represents mean ± SEM, based on 13-16 cells per condition. (**D**) Quantification of peak nuclear centroid shift immediately following nuclear ablation. Red line represents mean, black error bars represent standard deviation. *, *p* < 0.05; **, *p* < 0.01 based on unpaired non-parametric Kruskal Wallis test. (**E**) Cross-sectional area of blebs formed after cortex ablation. *, *p* < 0.05 for Friedman test; ****, *p* < 0.0001 based on unpaired non-parametric Kruskal-Wallis test. Cell counts per condition shown below graph. Red lines represent mean, black error bars represent standard deviation.

Targets for laser ablation were selected based on previously hypothesized roles in applying pushing or pulling forces on the nucleus during nuclear transit, including: *(i)* perinuclear actin filaments at the leading edge of the nucleus (“Nuclear Front”), which pull the nucleus forward, *(ii)* the rear cortex, whose contraction pushes the nucleus from behind (“Rear”), *(iii)* the front cortex at the leading edge of the cell, which may retain pressure in the front compartment of the cell that could inhibit nuclear transit (“Front”), *(iv)* both cortexes simultaneously to compare compartmentalized pressure in the front and rear of the cell body (“Both”), and *(v)* non-ablated cells to control for spontaneous movement of the nucleus during experiments (**Fig. 4A**). If contraction of the rear cortex pushes the nucleus forward, then ablation of the rear cortex during nuclear transit is expected to result in the nucleus falling backwards (**Supp. Fig. S5C**). On the other hand, if the nucleus is pulled forward by actomyosin fibers at the leading edge of the nucleus, as has been previously shown in MEFs (Davidson et al., 2020), then ablation at the nuclear front should result in the nucleus moving backwards (**Supp. Fig. S5C**). In contrast, ablation of the front cortex, which may inhibit forward movement of the nucleus through maintaining increased pressure in the front compartment of the cell (Petrie et al. 2014), is expected to result in forward movement of the nucleus (**Supp. Fig. S5C**). In all cases, movement of the nucleus in response to laser ablation is expected to occur almost instantaneously, as ablation should immediately disrupt the balance of forces acting on the nucleus, and nuclear movement will only be resisted by viscous forces within the cytoplasm (in the case of rearward movement) or the resistance of microfluidic constrictions (in the case of forward movement).

As seen in previous studies (Charras et al., 2005; Mistriotis et al., 2019; Tinevez et al., 2009), ablation of the cell cortex caused the formation of a large plasma membrane bleb, indicating the release of cytosolic pressure (**Fig. 4B, Arrow**). Ablation of the rear cortex and formation of the bleb at the cell posterior led to transient rearward movement of the cell nucleus (**Fig. 4B-D, Supp. Fig. S5D**). This observation supports our hypothesis that contraction of the rear cortex pushes the nucleus from behind by increasing intracellular pressure in the rear compartment. Furthermore, recovery and contraction of the bleb within a minute of the initial laser ablation resulted in forward movement of the nucleus (**Fig. 4B-D**), indicating that restoration of the cell rear cortex increased intracellular pressure in the back of the cell and pushed the nucleus forward.

Following ablation of the front cortex of cells, the nucleus immediately moved forwards, suggesting that prior to ablation, pressure in the front compartment of the cell impeded forward movement of the nucleus (**Fig. 4C-D, Supp. Fig. S5A, Supp. Fig. S5D**). However, the magnitude of the nuclear forward movement was less than the rearward movement observed when ablating the rear cortex, suggesting that pressure in the rear compartment may be greater than the pressure in the front compartment (**Fig. 4C-D**). To assess directly whether the pressure is greater in the rear compartment of the cell than in the front compartment, we ablated both front and rear cortexes of the cell simultaneously (**Fig. 4C-D, Supp. Fig. S5D**). Ablation of both cortexes led to a rearward shift of the nucleus, confirming that the pushing force generated by the pressure from the rear cortex contraction was greater than the resisting force generated by pressure from the front compartment (**Fig. 4C-D**).

As an additional means of comparing pressure between the front and rear compartments of the cell, we measured the size of blebs in the cell cortex formed as a result of cortex ablation (**Fig. 4E**). Previous work had demonstrated that bleb size following laser ablation increases as a consequence of elevated intracellular pressure and is limited only by the elasticity of the cell membrane (Tinevez et al., 2009). Thus, increased bleb sizes following laser ablation indicate increased intracellular pressure prior to ablation. Following simultaneous ablation of the front and rear cortexes, blebs formed in the rear compartment were significantly larger than blebs in the front compartment, indicating that pressure in the rear of the cell is greater than pressure in the front compartment (**Fig. 4E**). Collectively, these findings confirm that increased pressure at the cell rear helps to push the nucleus forward through constrictions.

Ablation directly in front of the nucleus led to a subtle rearward shift of the nucleus, suggesting the presence of pulling forces at the leading edge of the nucleus (**Fig. 4 B-C, Supp. Fig. S5A, Supp. Fig. S5D**), consistent with previous findings (Davidson et al., 2020). However, the rearward movement of the nucleus was very modest in comparison to the movement following rear cortex ablation (**Fig. 4D**). These findings indicate that the cortex-driven cells may rely more heavily on pushing forces from behind the nucleus, than on pulling forces at the nuclear front, which is consistent with the faster nuclear transit we observed in the cells using the cortex-driven mechanism (**Fig. 1D**).

We hypothesized that these intracellular pressure-driven pushing forces resulted from the accumulation and contraction of actomyosin in the rear cortex during cortex-driven nuclear transit. As we observed that ROCK-inhibition reduced myosin accumulation in the rear cortex, we hypothesized that ROCK inhibition reduces intracellular pressure behind the nucleus. Following Y27632 treatment, nuclear movement following rear cortex ablation was reduced, whereas nuclear shift in response to ablation of the nuclear front was unaffected (**Fig. 4D**), indicating that pushing forces between the rear cortex and nucleus result from ROCK-dependent actomyosin contractility. Additionally, ROCK inhibition led to a substantial decrease in the size of blebs formed following laser ablation (**Fig. 4E**), further confirming that ROCK mediated actomyosin contractility drives contraction of the rear cortex and increased cytosolic pressure in the rear compartment of the cell.

## Discussion

In this study, we investigated the cytoskeletal dynamics and intracellular forces acting on the nucleus as cells migrate through confined spaces. We identified a mechanism by which contraction of actomyosin at the rear cortex pushes the nucleus through constrictions during 3D migration. Our results suggest that cortex-driven nuclear transit is a distinct mechanism from alternative “non-cortex-driven” mechanisms, which have been described previously (Davidson et al., 2020; Petrie et al., 2014). In the cortex-driven nuclear transit mechanism, pushing forces at the cell rear are generated through contraction of actomyosin in the posterior cortex, which elevates cytosolic pressure behind the nucleus to push the nucleus forwards (**Fig. 5**). This contrasts the “non-cortex-driven” modes, in which pulling forces, through actomyosin or intermediate filaments, are mediated through physical connections between the nucleus and cytoskeleton (Davidson et al., 2020; Petrie et al., 2014; Thomas et al., 2015) (**Fig. 5**). Our studies suggest that some cell lines can use both mechanisms to achieve nuclear transit, though individual cell lines may have a propensity to use one mechanism over another. It remains to be elucidated whether these pushing and pulling mechanisms work independently to accomplish nuclear transit, or whether they act complimentarily.

**Figure 5:**
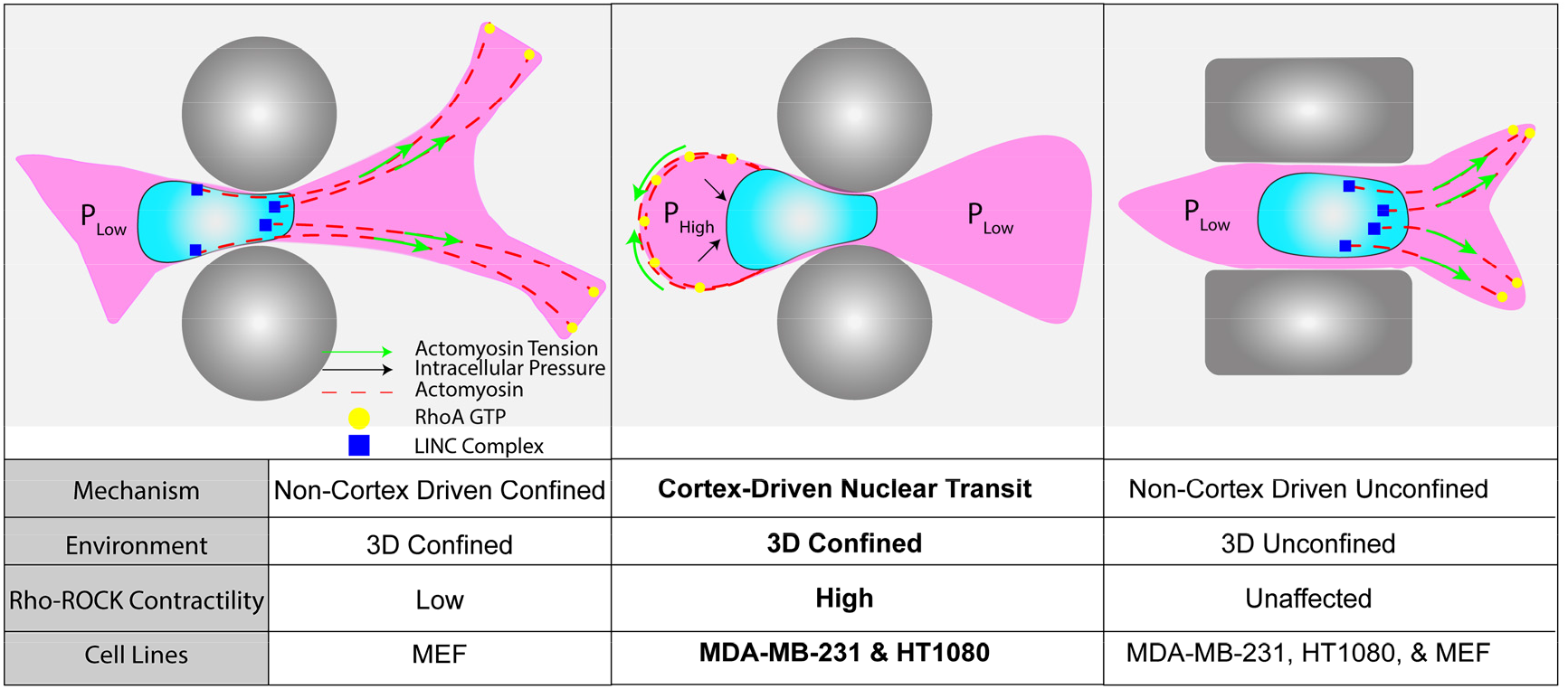
Model of alternate mechanisms of nuclear transit through constrictions. Pushing forces are generated through contraction of the posterior cortex, caused by co-localization of active RhoA, Myosin II, and Actin. Co-localization of these same proteins at the leading edge of the nucleus can generate pulling forces at the front of the nucleus, mediated through the LINC complex.

Mirroring our findings in microfluidic devices and collagen matrices, accumulation of actomyosin in the rear of cells migrating in 3D has been observed in MDA-MB-231 cells invading through Matrigel (Poincloux et al., 2011) and in vitro models of transendothelial migration (Chen et al., 2016), indicating that rear cortex-driven nuclear transit may play an important role in these physiological processes. Earlier studies of 3D dendritic cell migration reported that retraction of the cell rear preceded nuclear squeezing through constrictions (Lämmermann et al., 2008). These studies showed that inhibition of contractility through treatment with blebbistatin and Y27632 delayed contraction of the cell posterior, which corresponded with a reduced migration rate in confined environments (Lämmermann et al., 2008; Schaar and McConnell, 2005). However, it was unclear whether this retraction represented a direct transmission of force between the rear cortex and the nucleus, or whether rear retraction followed the forward movement of the nucleus. More recent findings have found elevated RhoA activity and myosin at the rear of cells migrating through 3D channels and showed evidence that this resulted in increased intracellular pressure behind the nucleus (Mistriotis et al., 2019). However, these studies did not address how intracellular pressure could contribute to nuclear transit through constrictions. Our results build upon these prior findings by confirming that retraction of the rear cortex pushes the nucleus forward during nuclear transit, and by demonstrating a functional role for cortex-driven pressure generation in confined 3D migration.

Our findings complement prior studies that have probed intracellular forces acting on the nucleus during cell migration. One such set of studies evaluated the role of pulling forces at the front of the nucleus, which increase intracellular pressure in the front compartment of fibroblasts and HT1080 cells migrating through 3D environments (Petrie et al., 2016, 2014). In contrast to our work, these studies used a probe to directly measure intracellular pressure in cells migrating through 3D cell-derived matrix. They found that actomyosin and intermediate filaments at the nuclear front pulled the nucleus forward to increase pressure at the leading edge of cells, which generated “lobopodia” to enable invasion through 3D environments. The apparent differences in the results may be due to the different cell types use or reflect differences in the geometry and physical properties of the microenvironment. For example, subsequent studies in mesenchymal stem cells migrating in 3D alginate hydrogels have indicated that pulling forces at the leading edge of the nucleus are not necessary for the formation of lobopodia (Lee et al., 2021). In the absence of nesprins, the nucleus was still able to move forward into constrictions to pressurize the front compartment of cells, while the authors noted an accumulation of pMLC in the rear cortex. These results mirror our findings of active RhoA in the rear cortex during nuclear entry to constrictions, supporting the idea that elevated pressure at the cell rear is increased at the beginning of nuclear transit.

One limitation of our studies is that our detection of intracellular forces must be inferred from protein localization and changes in nuclear position following laser ablation, rather than measuring these forces directly, as the PDMS enclosure of the microfluidic migration devices prevents direct access to the migrating cells. Our laser ablation assay is limited as an indirect measurement of intracellular pressure (or other cytoskeletal forces), as it only reveals forces acting on the nucleus at a single point in time, though the pressure throughout the cell body during nuclear transit may be highly dynamic as the nucleus shifts from the rear compartment of the cell to the front compartment. Additionally, the laser ablation likely disrupts not only actomyosin but also other force-generating cytoskeletal structures within the target region, including intermediate filaments and microtubules, which could further contribute to the force transmission and force generation. Nonetheless, as we found that the cortex-driven mechanism strongly depends on the Rho-ROCK-myosin pathway, we believe that this mechanism principally depends on actomyosin activity, though our results cannot rule out potential contributions of other cytoskeletal elements, which have previously been implicated in nuclear movement (Petrie et al., 2014; Tsai et al., 2010). In addition, the distinct circular shape of rear cortex, in contrast with that of front cortex, further supports the conclusion that the pressure is higher in the rear compartment.

A recent report demonstrated the role of pulling forces in mediating nuclear transit in MEFs (Davidson et al., 2020). Similar to our results in MDA-MB-231 cells, the authors demonstrated that ROCK inhibition increases nuclear transit times of MEFs migrating through narrow constrictions, but not through unconfined 3D channels (Davidson et al., 2020). Using a laser ablation assay, as we have used here, they found that ablation in front of the nucleus caused a rearward shift in the nucleus whereas ablation at the cell rear had no effect on nuclear movement. They also found that the rearward shift in nuclear position following ablation at the nuclear front was dependent on connections between the nucleus and cytoskeleton. These results contrast with our data in MDA-MB-231 cells, as we detected only very subtle nuclear shifts following ablation of the nuclear front, a significant rearward shift of the nucleus after rear cortex ablation, and LINC complex disruption did not impede nuclear transit (**Supp. Fig. S3**). This difference in results could be attributable to our findings that MEFs rarely use the cortex-driven nuclear transit mechanism in comparison to MDA-MB-231s, which very frequently used the cortex-driven mode. These two findings are complementary in that they highlight a key difference between MEFs, which in the “non-cortex-driven” mode apply pulling forces at the front of the nucleus (Davidson et al., 2020), and cortex-driven MDA-MB-231 cells which push the nucleus from behind through intracellular pressure generated by the rear cortex.

In our studies using a dominant negative nesprin construct (DN-KASH), LINC complex disruption had context-dependent effects on nuclear transit time. In MDA-MB-231 cells, DN-KASH expression did not alter nuclear transit times in cells using rear cortex driven migration but increased nuclear transit times in cells using non-cortex drive migration, suggesting that the LINC complex is important for pulling the nucleus forward. Surprisingly, DN-KASH expression did not significantly alter nuclear transit times in MEFs, suggesting cell-type specific or context dependent differences.

Although we have noted a cell-type specific preference towards the cortex-driven mechanism, and that the mechanism depends on high Rho-ROCK-mediated contractility, it remains unclear which biological and/or environmental factors cause individual cells to adopt a particular nuclear transit mechanism. As others have found that pulling forces at the front of the nucleus depend on the LINC complex (Davidson et al., 2014; Petrie et al., 2016), it is possible that the cortex-driven mechanism could compensate for a lack of pulling forces in specific scenarios. For example, pulling forces at the leading edge of the cell require adhesions to the surrounding environment. Thus, cells that migrate in an adhesion-independent “amoeboid” migration mode may require cortex-driven forces for nuclear transit through narrow constrictions, or cells may benefit from an additional ‘boost’ from the cortex-driven pressure when pulling forces are insufficient to move the nucleus through a constriction. Other studies have indicated that deformation of the nucleus or collapse of the microtubule network in the rear compartment during confined migration may actively trigger contractility at the cell rear (Ju et al., 2023; Lomakin et al., 2020). Our work supplements these findings by demonstrating how contraction of the cell rear may push the nucleus forward to pass through constrictions. Future studies will be necessary to determine the specific factors determining the selection of a particular migration mechanism, and whether this is an active decision-making process requiring specific signaling pathways, or whether cells use multiple migration modes in parallel, with a specific mode dominating depending on the physical microenvironment (Friedl and Wolf, 2010; Geum et al., 2016; Liu et al., 2015; Wolf et al., 2013).

In conclusion, the recognition that myosin II and Rho/ROCK mediated rear cortex contraction aids in nuclear transit adds new insights into the mechanism by which cells traverse confining 3D environments, and further highlights the plasticity of cells to adapt their migration mode to the specific conditions. An improved understanding of the diverse mechanisms cells have available to overcome narrow interstitial spaces, and how cells select between these mechanisms, may lead to new targets for therapies to reduce or prevent cancer cell invasion and metastasis, or to boost the migration of immune cells to sites of infection.

## Methods

### Cell Culture

MDA-MB-231 metastatic breast adenocarcinoma cells were purchased from American Type Culture Collection (ATCC); HT1080 fibrosarcoma cells were a gift from Peter Friedl and Katarina Wolf, originally purchased from DSMZ in Braunschweig, Germany; MEFs expressing GFP-Nesprin-2 were a gift from Cecile Sykes; MDA-MB-231 cells expressing the RhoA-FRET biosensor were a gift from Louis Hodgson. All cell lines were cultured in Dulbecco’s Modified Eagle Medium (DMEM) supplemented with 10% (v/v) fetal bovine serum (FBS, Seradigm VWR), and penicillin and streptomycin (50 U/mL, ThermoFisher Scientific) and were maintained at 37°C and 5% CO2. Cell lines were tested for mycoplasma infection following completion of experiments. Human cell lines were verified through the ATCC Cell Line Authentication Service.

### Treatment with Rho/ROCK inhibitors and activators

For RhoA inhibition/activation experiments, cells were treated with Y-27632 (10 µM; #1000583 Cayman Chemical), LPA (10 µM; #L7260 Sigma Aldrich), RhoA Activator II (1 µg/mL, #CN03 Cytoskeleton, Inc.), or a vehicle (DMSO) control.

### Viral and Piggybac Modification

Cell lines were stably modified to express fluorescently tagged proteins using lentiviral vectors (pCDH-mCherry2-Actin Chromobody-puro and pCDH-mNeonGreen-H2B Histone-puro). Transient expression of CMV-GFP-NMHC-IIA (Addgene #11347) and mCherry-MyosinIIB-N-18 (Addgene #55107) was achieved using a Nucleofector II electroporator and the Cell Line Nucleofector Kit V (Lonza Biosciences). MEFs and MDA-MB-231 cells were stably modified using BFP and DN-KASH expression plasmids in a doxycycline-inducible Piggybac plasmid backbone (Addgene #187019). For lentiviral modifications, pseudoviral particles were produced using 293TN cells (System Biosciences, SBI). The 293TN cells were transfected with the lentiviral plasmid of interest as well as lentiviral packaging and envelope plasmids (psPAX and pMD2.G, gifts from Didier Trono) using PureFection transfection reagent (SBI), per manufacturer’s protocol. Supernatants were collected at 48-, 72-, and 96-hour intervals post-transfection and passed through a 0.45 µm filter to remove cells and cellular debris. Cells to be transduced were cultured for 24 hours prior to transduction at a confluency of 50% and were then cultured in lentivirus-containing media with polybrene (8 µg/mL). Lentivirus containing media was removed after 24 hours, and following an additional 24 hours of recovery time, cells were selected using 1 µg/mL of puromycin (Invivogen) for 3 days. For Piggybac modifications, MEFs were seeded into 6-well plates and were transfected with 1.75 µg of Piggybac plasmids and 0.75 µg of a hyperactive transposase using PureFection Transfection reagent (SBI), per manufacturer’s protocol. Transfected cells were selected using 800 µg/mL G418 for 3 days. Cells were sorted using flow assisted cell sorting (FACS) in order to obtain consistent expression levels.

### LINC complex disruption and confirmation of nesprin displacement in MEFs and MDA-MB-231 cells

MEFs with fluorescently labeled endogenous Nesprin2 (GFP-Nesprin-2, Davidson et al. 2020), and expressing an mCherry2-Actin chromobody and either the inducible BFP-DN-KASH or BFP-only control construct were treated with 1 µg/mL doxycycline for 24 hours. Cells were subsequently fixed with 4% PFA for 15 minutes, washed with 1× PBS, and mounted on glass slides using Hydromount (Electron Microscopy Sciences).

For LINC complex disruption in MDA-MB-231 cells, cells expressing either the inducible BFP-DN-KASH or BFP-only control construct were treated with 1 µg/mL doxycycline for 24 hours. Cells were subsequently fixed with 1:1 methanol:acetone at –20°C for 15 minutes and washed with 1× PBS. Cells were then washed with 1 × PBS and blocked in 3% BSA with 0.1% Triton-X 100 (ThermoFisher Scientific) and 0.1% Tween (Sigma) in PBS for 1 hour at room temperature. To confirm the displacement of endogenous nesprins, cells were immunofluorescently labeled with an antibody directed against human Nesprin-1 (Developmental Studies Hybridoma Bank, MANNES1E(8C3), supernatant form) at a 1:3 dilution used overnight at 4°C. The following day, the cells were washed 3-times with 1× PBS and stained with Alexa Fluor 568 goat anti-mouse IgG1 secondary antibody (Life Technologies, catalog # A21124) at room temperature for one hour. Coverslips were washed 3 times with 1X PBS and mounted on glass slides using Hydromount (Electron Microscopy Sciences).

Nesprin displacement was measured manually through observation of the GFP-Nesprin-2 signal using the ImageJ/FIJI software. Cells with high GFP-Nesprin-2 signal around the periphery of the nucleus were defined as “non-displaced”, whereas cells lacking this enrichment of GFP-Nesprin-2 at the nuclear envelope were defined as “displaced” (see Supp. Fig. 3A and 3E). For doxycycline-treated cells, only BFP-positive cells were included for analysis. For cells that were not treated with doxycycline, all cells within the field of view were included in the analysis, since BFP expression requires doxycycline treatment. All analyses were conducted with the observer blinded to the test groups during the determination of Nesprin displacement.

### Microfluidic Device Preparation

Microfluidic devices were prepared as described previously (Davidson et al., 2015; Keys et al., 2018). Briefly, microfluidic devices were made through soft lithography by curing polydimethylsiloxane (PDMS) in a plastic mold containing microfluidic features at 65°C for 2 hours. After removing cured PDMS from the mold, PDMS features were cut out using biopsy punches, and then both PDMS devices and glass coverslips were washed with deionized water and isopropyl alcohol, and then treated in a plasma cleaner (Harrick Plasma) for 5 minutes. PDMS pieces were then bound to activated coverslips and then heated at 95°C on a hotplate for 1 minute to improve adhesion. After heating, devices were filled with 70% ethanol to sterilize internal surfaces, rinsed with PBS, and then coated with extracellular matrix proteins. For MDA-MB-231 cells, devices were coated with 50 µg/mL type-I rat tail collagen (Corning) in 0.02 N glacial acetic acid overnight at 4°C. For MEFs and HT1080 cells, devices were coated with 50 µg/mL fibronectin (Millipore) in PBS overnight at 4°C. Before seeding with cells, devices were rinsed once with PBS and once with cell media. Devices were then aspirated of all media, and then loaded with cell-containing solution (roughly 20,000-30,000 cells per chamber). In devices, cells were cultured in Fluorobrite DMEM supplemented with 10% FBS (Seradigm, VWR), penn/strep (50 U/mL, ThermoFisher), GlutaMAX, and 10 mM HEPES (Gibco). Cells were allowed to attach within devices for 8-12 hours before live cell imaging. Cells were imaged for 12-14 hours at 3-to-5-minute intervals on a Zeiss LSM700 laser scanning confocal microscope at 37°C. For migration experiments with MEFs and MDA-MB-231s expressing either the inducible BFP-DN-KASH or BFP-only control constructs, the BFP channel was acquired using the 405-nm laser excitation line only at the beginning and end of each experiment to confirm BFP expression while minimizing phototoxic effects. Only cells expressing BFP-DN-KASH or BFP-only were used in the migration analysis.

### Analysis of Migration Mode and Nuclear Transit Times

Measurements of nuclear transit times were performed as described previously (Davidson et al., 2015). Briefly, cells were imaged at 5-minute intervals on a Zeiss LSM710 laser scanning confocal microscope using 488-nm and 561-nm laser lines as they migrated through migration devices over a period of 12-16 hours. A fluorescent marker of the nucleus (either mNeonGreen-H2B Histone, or GFP-Nesprin2) was used to track the position of the nucleus relative to microfluidic constrictions. An imaginary line was marked 5 microns on either side of the middle of the constriction, to define when the nucleus “enters” and “exits” the constriction. The times at which the nucleus enters and passes through the constriction were noted, and this interval was recorded as the nuclear transit time. Cells which did not complete nuclear transit following entry to the constriction were excluded from analysis. These criteria were established prior to performing experiments, based on our group’s prior work with nuclear transit experiments.

### Quantification of actin and myosin II localization

Quantification of fluorescent protein localization during nuclear transit was performed manually in the ImageJ/FIJI software package. The region of interest (i.e., rear cortex (RC), front cortex (FC), nuclear front (NF), or whole cell) was manually traced in the fluorescent channel of interest, and the average intensity of the fluorescent signal was measured (See Supp. Fig. 1). The rear cortex (RC) region was defined as a 2 µm wide region starting at the rear perimeter of the cell, behind the nuclear constriction area. The front cortex (FC) region was defined as a 2 µm wide region starting at the front perimeter of the cell, in front of the nuclear constriction area. The nuclear front (NF) region was defined as a 2 µm wide region starting at the leading edge of the nucleus. The fluorescent signal was measured at a single time point during the specified stage of nuclear transit (i.e. “Pre”, “Entry”, “Exit”, “Post).

### Collagen Matrix Cell Migration Assay

Collagen matrix cell migration assays were performed as described previously (Cross et al., 2010). In brief, PDMS wells were formed by punching out 10 mm diameter holes from PDMS slices. PDMS and glass coverslips were treated in a plasma cleaner for 5 minutes, before bonding PDMS to treated glass slides on a 95°C hot plate for 3 minutes. Wells were coated with 1% PIE for 10 minutes, followed by 0.1% Glutaraldehyde for 30 minutes, and then subsequently washed with PBS once, and cell media once, before adding the collagen solution. Collagen matrices were prepared by mixing Type I rat tail collagen (Corning) with DMEM and NaOH (to reach a neutral pH of 7.4) with a solution of cells suspended in complete DMEM to achieve a final density of 80,000 cells/mL. Small volumes of collagen solution (10-15 µL) were pipetted into the middle of wells, where they were allowed to polymerize at 37°C for 30 minutes. Following polymerization, wells were filled with complete DMEM and were subsequently incubated at 37°C for 48 hours before imaging experiments.

### RhoA FRET Imaging and Analysis

Analysis of RhoA-FRET biosensors was performed as described previously (Cheung et al., 2022). Cells were imaged at regular intervals on a dual channel image acquisition set-up to minimize temporal and spatial misalignment problems which impair FRET result interpretation. Cells were imaged at 5-minute intervals for 12 to 16 hours. Cells were excited with a ET436/20× (CFP Excitation) source for an exposure time of 3 seconds with CFP excitation while CFP and YFP (“FRET”) emission channels were captured simultaneously, by using a W-View Gemini beam splitter (A12801-01, Hamamatsu Photonics) and a T505lpxr dichroic mirror, through ET480/40m and ET535/30m optical filters, respectively. The light source for fluorescence imaging was a xenon arc lamp (Lambda LB-LS/30, Sutter Instruments). An absorptive neutral density filter with OD = 0.3 (NE03B-A, ThorLabs) was used to attenuate light to minimize phototoxicity. All filters and dichroic mirrors were purchased from Chroma Technology. Prior to each experiment, the x-y positions of CFP and FRET channels were adjusted using the W-VIEW Adjustment software (Hamamatsu Photonics) to achieve a reasonable alignment for easier image registration and processing. The epifluorescence microscope was surrounded by an incubator with a temperature of 37°C, 5% CO_2_, and humidity of ∼70%.

Field alignment, flatfield correction, and background subtraction were performed as described previously (Cheung et al., 2022). Images of cells were then manually masked by tracing the outline of cells using the CFP channel in ImageJ/Fiji. Following these corrections to both the CFP and FRET channels, FRET ratios were calculated by dividing the FRET channel by the CFP channel. The sensitivity of the RhoA-FRET biosensor expressed in this MDA-MB-231 cell line to RhoA activation and inactivation was performed under identical experimental conditions on this imaging set-up as described previously (Cheung et al., 2022). Local FRET ratios were then calculated by manually tracing regions as defined in Supp. Fig. S4A.

### Laser Ablation of Actin

Cells were seeded into migration devices 12-16 hours before ablation experiments. Cells were maintained at 37°C in Fluorobrite DMEM containing 10% FBS, 10 mM HEPES (Gibco), GlutaMax, and penicillin and streptomycin (50 U/mL). Cells were imaged through a water immersion 40×/NA 1.1 objective (Carl Zeiss Imaging) using 488-nm and 561-nm laser lines. Cells were imaged at 3 second intervals for 63 frames. After 9 seconds, cells were ablated in defined regions using a 780 nm laser produced by a Spectra Physics Insight multiphoton excitation source. Ablation was performed at 40-70% attenuation, following ablation tests at the beginning of each experiment to verify that the laser successfully disrupted the actin cytoskeleton without causing cell death. Cells which exited the constriction, displayed rapid blebbing, or appeared to die following laser ablation were excluded from analysis, while cells which were able to recover following disruption of the actin network were included. Exclusion criteria were established prior to conducting experiments, based on prior experience with laser ablation assays (Mistriotis et al., 2019).

### Image Analysis of Laser Ablation Experiments

Tracking of nuclear movement following laser ablation was achieved using ImageJ/FIJI software and a custom MATLAB script. Briefly, the image channel containing the fluorescent mNeonGreen-H2B Histone signal was processed using a 3-pixel median filter to eliminate noise, and then a binary mask of this signal was generated using ImageJ/FIJI’s built-in RenyiEntropy masking function. This binary mask was then processed in MATLAB to track the position of the nucleus’ centroid at each time point. The furthest position of nuclear movement prior to recovery (within 20 seconds of the initial ablation) was considered as the direct result of force disruption as a consequence of laser ablation and was recorded as the peak centroid shift (**Supp. Fig. S5E**). Bleb area measurements were performed in ImageJ/FIJI at the time point when the bleb had reached its maximum size. Outer bleb perimeter was traced manually, and inner bleb perimeter was traced at the periphery of the cell at the time point immediately prior to laser ablation.

### Statistical Analysis

All experimental data are based on at least three independent experiments, unless otherwise specified. Statistics tests were performed, and graphs were created using GraphPad Prism (GraphPad Software, San Diego, USA). Nuclear transit times were compared using a one-way ANOVA. Frequency of nuclear transit mechanism selection and nesprin displacement analysis was performed using a Chi Squared test of independence. Changes in local FRET ratio, actin accumulation, and myosin accumulation during nuclear transit were evaluated using a paired non-parametric ANOVA test (Friedman). Nuclear centroid shifts following laser ablation were compared using an unpaired non-parametric ANOVA test (Kruskal-Wallis). Bleb formation in cells ablated at a single site were compared using an unpaired non-parametric ANOVA test (Kruskal-Wallis) while bleb formation in cells ablated at both cortexes simultaneously were compared using a paired non-parametric ANOVA test (Friedman).

## Acknowledgements

We thank Louis Hodgson for providing the MDA-MB-231 cell line expressing the RhoA-FRET Biosensor and Peter Friedl and Katarina Wolf for providing the HT1080 fibrosarcoma cell line. The mouse embryonic fibroblast cell line expressing Nesprin-2 GFP was made by Aude Battistella and Patricia Davidson and was a kind gift from Cécile Sykes and Sirine Amiri. We thank Klaus Hahn and Louis Hodgson for their advice regarding the use and analysis of FRET biosensors in migrating cells. We thank Philipp Isermann for his help in setting up the initial laser ablation experiments on MDA-MB-231 cells.

## References

Bone, C.R., Chang, Y.T., Cain, N.E., Murphy, S.P., Starr, D.A., 2016. Development 143, 4193–4202.

Caille, N., Thoumine, O., Tardy, Y., Meister, J.J., 2002. J Biomech 35, 177–187.

Charras, G.T., Yarrow, J.C., Horton, M.A., Mahadevan, L., Mitchison, T.J., 2005. Nature 435, 365–369.

Chen, M.B., Lamar, J.M., Li, R., Hynes, R.O., Kamm, R.D., 2016. Cancer Res 76, 2513–2524.

Cheung, B.C.H., Hodgson, L., Segall, J.E., Wu, M., 2022. Exp Cell Res 410, 112939.

Cross, V.L., Zheng, Y., Won Choi, N., Verbridge, S.S., Sutermaster, B.A., Bonassar, L.J., Fischbach, C., Stroock, A.D., 2010. Biomaterials 31, 8596–8607.

Davidson, P.M., Battistella, A., Déjardin, T., Betz, T., Plastino, J., Borghi, N., Cadot, B., Sykes, C., 2020. EMBO Rep 21, e49910.

Davidson, P.M., Denais, C., Bakshi, M.C., Lammerding, J., 2014. Cell Mol Bioeng 7, 293–306.

Davidson, P.M., Sliz, J., Isermann, P., Denais, C.M., Lammerding, J., 2015. Integr Biol 7, 1534–1546.

Denais, C.M., Gilbert, R.M., Isermann, P., McGregor, A.L., te Lindert, M., Weigelin, B., Davidson, P.M., Friedl, P., Wolf, K., Lammerding, J., 2016. Science (1979) 352, 353–358.

Doyle, A.D., Petrie, R.J., Kutys, M.L., Yamada, K.M., 2013. Curr Op Cell Bio 25, 642–649.

Friedl, P., Wolf, K., 2010. J Cell Biol 188, 11–9.

Gadea, G., De Toledo, M., Anguille, C., Roux, P., 2007. Journal of Cell Biology 178, 23–30.

Geiger, F., Rüdiger, D., Zahler, S., Engelke, H., 2019. PLoS One 14.

Geum, D.T., Kim, B.J., Chang, A.E., Hall, M.S., Wu, M., 2016. Eur Phys J Plus 131.

Harada, T., Swift, J., Irianto, J., Shin, J.-W., Spinler, K.R., Athirasala, A., Diegmiller, R., Dingal, P.C.D.P., Ivanovska, I.L., Discher, D.E., 2014. Journal of Cell Biology 204, 669–682.

Hetmanski, J.H.R., de Belly, H., Busnelli, I., Waring, T., Nair, R. v, Sokleva, V., Dobre, O., Cameron, A., Gauthier, N., Lamaze, C., Swift, J., del Campo, A., Starborg, T., Zech, T., Goetz, J.G., Paluch, E.K., Schwartz, J.-M., Caswell, P.T., 2019. Dev Cell 51, 460-475.e10.

Jayo, A., Malboubi, M., Antoku, S., Chang, W., Ortiz-Zapater, E., Groen, C., Pfisterer, K., Tootle, T., Charras, G., Gundersen, G.G., Parsons, M., 2016. Dev Cell 38, 371–383.

Ju, R.J., Falconer, A.D., Dean, K.M., Fiolka, R.P., Sester, D.P., Nobis, M., Timpson, P., Lomakin, A.J., Danuser, G., White, M.D., Oelz, D.B., Haass, N.K., Stehbens, S.J., 2023. bioRxiv 2022.02.08.479516.

Kalukula, Y., Stephens, A.D., Lammerding, J., Gabriele, S., 2022. Nat Rev Mol Cell Biol 23, 583–602.

Keys, J., Windsor, A., Lammerding, J., 2018. Assembly and Use of a Microfluidic Device to Study Cell Migration in Confined Environments, in: Gundersen, G.G., Worman, H.J. (Eds.), The LINC Complex: Methods and Protocols. Springer New York, New York, NY, pp. 101– 118.

Khatau, S.B., Bloom, R.J., Bajpai, S., Razafsky, D., Zang, S., Giri, A., Wu, P.-H., Marchand, J., Celedon, A., Hale, C.M., Sun, S.X., Hodzic, D., Wirtz, D., 2012. Sci Rep 2, 488.

Lammerding, J., 2011. Compr Physiol 1, 783–807.

Lämmermann, T., Bader, B.L., Monkley, S.J., Worbs, T., Wedlich-Söldner, R., Hirsch, K., Keller, M., Förster, R., Critchley, D.R., Fässler, R., Sixt, M., 2008. Nature 453, 51–55.

Lee, H., Alisafaei, F., Adebawale, K., Chang, J., Shenoy, V.B., Chaudhuri, O., 2021. Sci Adv 7, eabd4058.

Liu, Y.-J., Le Berre, M., Lautenschlaeger, F., Maiuri, P., Callan-Jones, A., Heuzé, M., Takaki, T., Voituriez, R., Piel, M., 2015. Cell 160, 659–672.

Lomakin, A.J., Cattin, C.J., Cuvelier, D., Alraies, Z., Molina, M., Nader, G.P.F., Srivastava, N., Saez, P.J., Garcia-Arcos, J.M., Zhitnyak, I.Y., Bhargava, A., Driscoll, M.K., Welf, E.S., Fiolka, R., Petrie, R.J., de Silva, N.S., González-Granado, J.M., Manel, N., Lennon-Duménil, A.M., Müller, D.J., Piel, M., 2020. Science (1979) 370.

Lombardi, M.L., Jaalouk, D.E., Shanahan, C.M., Burke, B., Roux, K.J., Lammerding, J., 2011. Journal of Biological Chemistry 286, 26743–26753.

Marks, P.C., Petrie, R.J., 2022. Phys Biol 19.

McGregor, A.L., Hsia, C., Lammerding, J., 2016. Curr Op Cell Bio 40, 32–40.

Mistriotis, P., Wisniewski, E.O., Bera, K., Keys, J., Li, Y., Tuntithavornwat, S., Law, R.A., Perez-Gonzalez, N.A., Erdogmus, E., Zhang, Y., Zhao, R., Sun, S.X., Kalab, P., Lammerding, J., Konstantopoulos, K., 2019. J Cell Biol 218, 4093–4111.

Pertz, O., Hodgson, L., Klemke, R.L., Hahn, K.M., 2006. Nature 440, 1069–1072.

Petrie, R.J., Harlin, H.M., Korsak, L.I.T., Yamada, K.M., 2016. Journal of Cell Biology 216, 93–100.

Petrie, R.J., Koo, H., Yamada, K.M., 2014. Science (1979) 342, 1062–1065.

Poincloux, R., Collin, O., Lizárraga, F., Romao, M., Debray, M., Piel, M., Chavrier, P., 2011. Proceedings of the National Academy of Sciences 108, 1943–1948.

Ridley, A.J., Schwartz, M.A., Burridge, K., Firtel, R.A., Ginsberg, M.H., Borisy, G., Thomas Parsons, J., Rick Horwitz, A., 2003. Cell Migration: Integrating Signals from Front to Back, Annu. Rev. Plant Physiol. Plant Mol. Biol. H. Hashimoto.

Rowat, A.C., Jaalouk, D.E., Zwerger, M., Ung, W.L., Eydelnant, I.A., Olins, D.E., Olins, A.L., Herrmann, H., Weitz, D.A., Lammerding, J., 2013. Journal of Biological Chemistry 288, 8610–8618.

Sahai, E., Marshall, C.J., 2003. Nat Cell Biol 5, 711–719.

Schaar, B.T., McConnell, S.K., 2005. Cytoskeletal coordination during neuronal migration, PNAS.

Thiam, H.R., Vargas, P., Carpi, N., Crespo, C.L., Raab, M., Terriac, E., King, M.C., Jacobelli, J., Alberts, A.S., Stradal, T., Lennon-Dumenil, A.M., Piel, M., 2016. Nat Commun 7.

Thomas, D.G., Yenepalli, A., Denais, C.M., Rape, A., Beach, J.R., Wang, Y.-L., Schiemann, W.P., Baskaran, H., Lammerding, J., Egelhoff, T.T., 2015. J Cell Biol 210, 583–594.

Tinevez, J.-Y., Schulze, U., Salbreux, G., Roensch, J., Joanny, J.-F., Paluch, E., 2009. Proceedings of the National Academy of Sciences 106, 18581–18586.

Tsai, J.W., Lian, W.N., Kemal, S., Kriegstein, A.R., Vallee, R.B., 2010. Nat Neurosci 13, 1463–1472.

Wang, H., Alarcón, C.N., Liu, B., Watson, F., Searles, S., Lee, C.K., Keys, J., Pi, W., Allen, D., Lammerding, J., Bui, J.D., Klemke, R.L., 2021. Nat Biomed Eng.

Weigelin, B., Bakker, G.-J., Friedl, P., 2012. Intravital 1, 32–43.

Wilson, M.H., Holzbaur, E.L.F., 2012. J Cell Sci 125, 4158–4169.

Wolf, K., te Lindert, M., Krause, M., Alexander, S., te Riet, J., Willis, A.L., Hoffman, R.M., Figdor, C.G., Weiss, S.J., Friedl, P., 2013. Journal of Cell Biology 201, 1069–1084.

Wolf, K., Mazo, I., Leung, H., Engelke, K., von Andrian, U.H., Deryugina, E.I., Strongin, A.Y., Bröcker, E.B., Friedl, P., 2003. Journal of Cell Biology 160, 267–277.

Wu, J., Lee, K.C., Dickinson, R.B., Lele, T.P., 2011. J Cell Physiol 226, 2666–2674.

Yadav, S.K., Feigelson, S.W., Roncato, F., Antman-Passig, M., Shefi, O., Lammerding, J., Alon, R., 2018. J Leukoc Biol 104, 239–251.

Yamada, K.M., Sixt, M., 2019. Nat Rev Mol Cell Biol 20, 738–752.

Zhu, R., Antoku, S., Gundersen, G.G., 2017. Current Biology 27, 3097–3110.e5.

