## Supplementary Materials for "Rear cortex contraction aids in nuclear transit during confined migration by increasing pressure in the cell posterior"

### Supplementary Figures

#### Figure S1

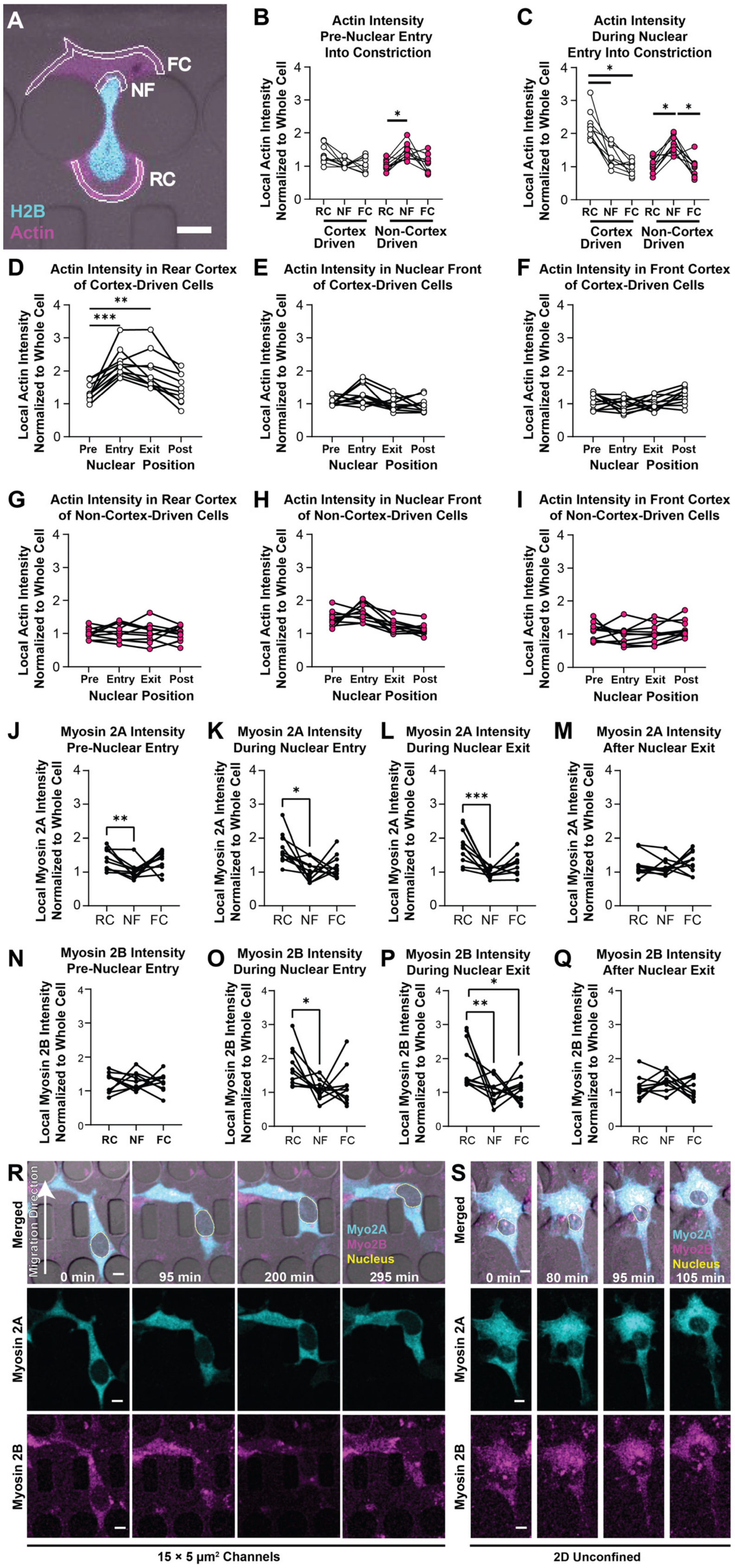

**Supplemental Figure 1. Quantification of localized actin and non-muscle myosin II accumulation in MDA-MB-231 cells during nuclear transit.** (**A**) Quantification strategy for local actin accumulation. The average intensity of actin signal in each white-outlined region was normalized to the average intensity of the whole cell body at each time point during nuclear transit. RC: Rear Cortex, FC: Front Cortex, NF: Nuclear Front. Scale bar: 10 μm (**B**) Actin intensity in specified cell regions prior to nuclear entry for cortex driven and non-cortex driven cells. (**C**) Actin intensity in specified cell regions during nuclear entry for cortex driven and non-cortex driven cells. (**D-F**) Time course of actin intensity in each specified region of cortex-driven cells during nuclear transit. (**G-I**) Time course of actin intensity in each specified region of non-cortex-driven cells during nuclear transit. Connected data points in each graph represent repeated measurements from the same cell. *, $p < 0.05$; **, $p < 0.01$; ***, $p < 0.001$, based on Friedman test. (**J-M**) Myosin IIA intensity normalized to average cell intensity in specified cell regions during specified time point in nuclear transit sequence. (**N-Q**) Myosin IIB intensity normalized to average cell intensity in specified cell regions during specified time point in nuclear transit sequence. Quantification process is identical to process illustrated in panel A. Connected data points in each graph represent repeated measurements from an individual cell. *, $p < 0.05$; **, $p < 0.01$; ***, $p < 0.001$, based on Friedman test. (**R-S**) Localization of myosin II to rear cortex is unique to MDA-MB-231 cells migrating through narrow constrictions and not observed during migration through 15×5 $\mu m^2$ control channels (R) or on collagen-coated 2D coverslip surface (S). Representative time lapse imaging series of MDA-MB-231 cells expressing GFP-myosin IIA and mCherry-myosin IIB. Scale bars: 10 μm.

**Figure S2**

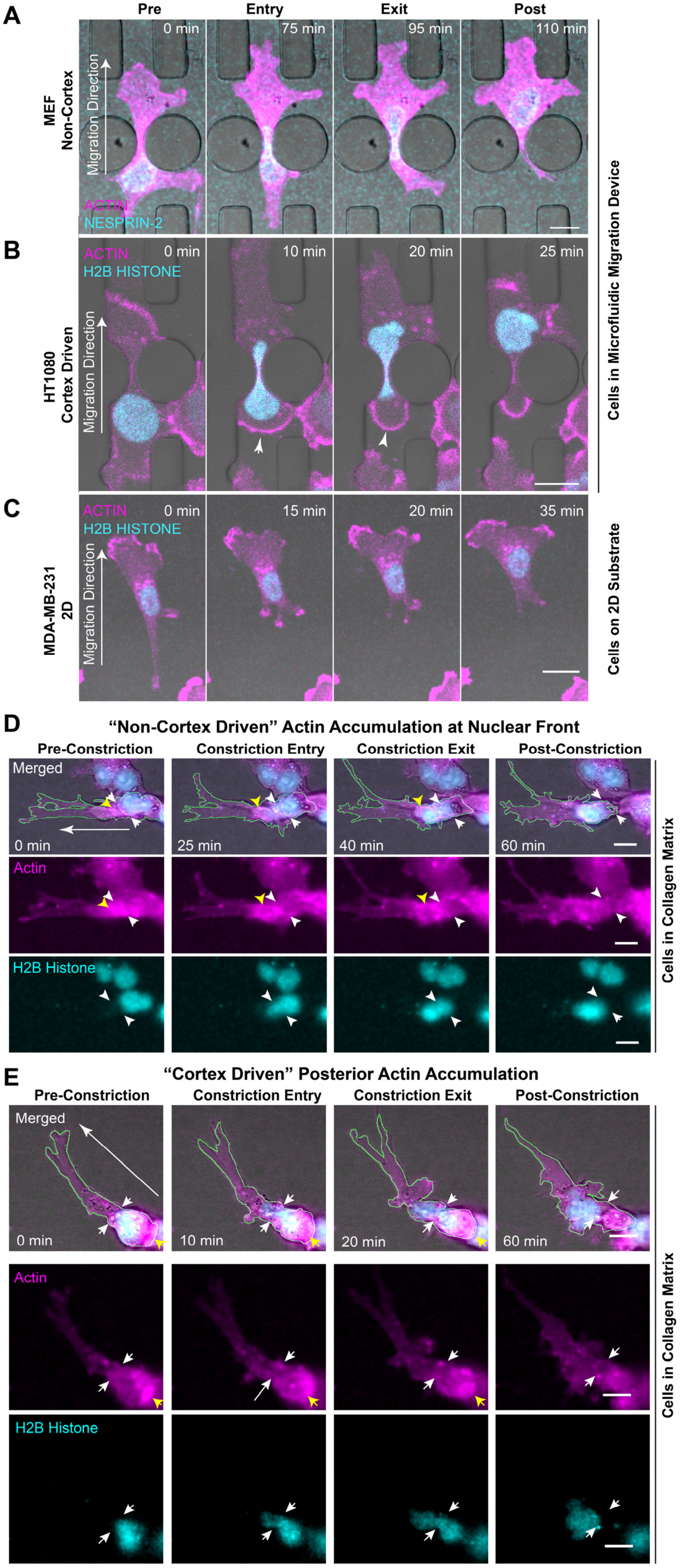
A
Pre
Entry
Exit
Post
0 min
75 min
95 min
110 min
MEF
Non-Cortex
Migration Direction
ACTIN
NESPRIN-2
B
ACTIN
H2B HISTONE
0 min
10 min
20 min
25 min
HT1080
Cortex Driven
Migration Direction
Cells in Microfluidic Migration Device
C
ACTIN
H2B HISTONE
0 min
15 min
20 min
35 min
MDA-MB-231
2D
Migration Direction
Cells on 2D Substrate
D
"Non-Cortex Driven" Actin Accumulation at Nuclear Front
Pre-Constriction
Constriction Entry
Constriction Exit
Post-Constriction
Merged
0 min
25 min
40 min
60 min
Actin
H2B Histone
Cells in Collagen Matrix
E
"Cortex Driven" Posterior Actin Accumulation
Pre-Constriction
Constriction Entry
Constriction Exit
Post-Constriction
Merged
0 min
10 min
20 min
60 min
Actin
H2B Histone
Cells in Collagen Matrix

**Supplementary Figure 2. Cells exhibit different nuclear transit mechanisms.** (**A-C**) Nuclear transit mechanisms depend on cell type and confinement. Representative time-lapse series of different cell morphologies observed during nuclear transit of MEFs (A) and HT1080 cells (B) in microfluidic devices, and image series of MDA-MB-231 cells migrating on a collagen-coated glass cover slip (C). When passing through narrow constrictions in the microfluidic migration devices, MEFs primarily use a non-cortex driven mechanism (A), in contrast to HT1080 cells, which use a cortex-driven mechanism (B; arrows). The cortex-driven mechanism used by MDA-MB-231 cells in Fig. 1B is unique to confined environments and is not observed during migration on collagen-coated 2D glass substrates (C). Scale bars: 20 µm. (**D-E**) MDA-MB-231 cells migrating in 3D collagen matrices exhibit at least two different modes of nuclear transit, reflected in distinct cytoskeletal actin organization and dynamics. In some cells, actin accumulates at the leading edge of the nucleus prior to and during nuclear transit (D). In other cells, actin accumulates at the cell posterior to generate "cortex-driven" nuclear transit as the rear cortex contracts (E). White arrow indicates direction of migration. Green outline in merged channel shows outline of cell body. Yellow arrowheads indicate actin accumulation. White arrowheads indicate apparent constriction within the collagen matrix. Scale bars: 10 µm. Magenta: mCherry2-Actin Chromobody. Cyan: mNeonGreen-H2B Histone.

Figure S3

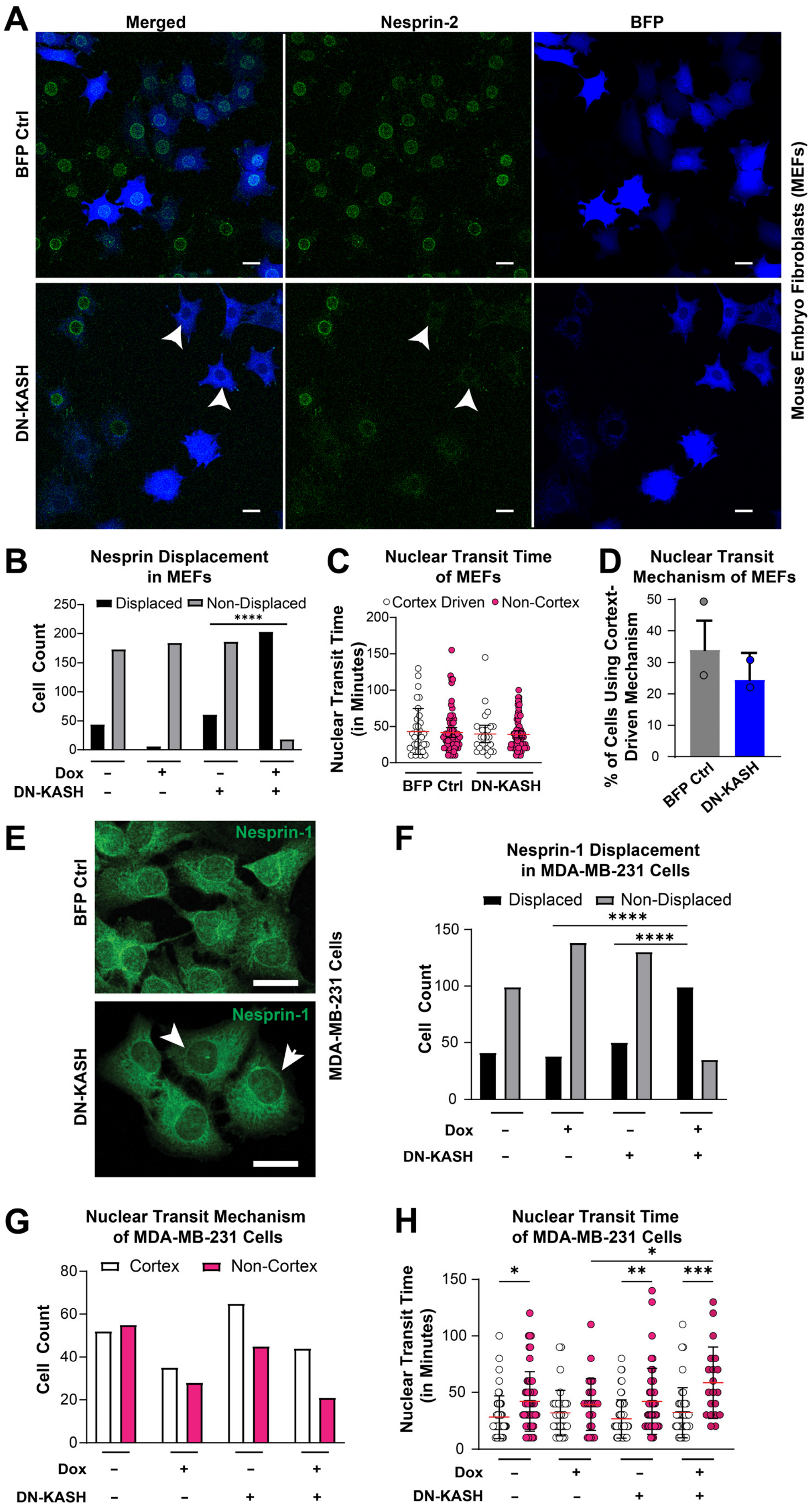
A
Merged
Nesprin-2
BFP
BFP Ctrl
DN-KASH
Mouse Embryo Fibroblasts (MEFs)
B
Nesprin Displacement in MEFs
Displaced
Non-Displaced
Cell Count
Dox
DN-KASH
C
Nuclear Transit Time of MEFs
Cortex Driven
Non-Cortex
Nuclear Transit Time (in Minutes)
BFP Ctrl
DN-KASH
D
Nuclear Transit Mechanism of MEFs
% of Cells Using Cortex-Driven Mechanism
BFP Ctrl
DN-KASH
E
Nesprin-1
BFP Ctrl
DN-KASH
MDA-MB-231 Cells
F
Nesprin-1 Displacement in MDA-MB-231 Cells
Displaced
Non-Displaced
Cell Count
Dox
DN-KASH
G
Nuclear Transit Mechanism of MDA-MB-231 Cells
Cortex
Non-Cortex
Cell Count
Dox
DN-KASH
H
Nuclear Transit Time of MDA-MB-231 Cells
Nuclear Transit Time (in Minutes)
Dox
DN-KASH

**Supplemental Figure 3. LINC Complex disruption has cell-type specific effects on confined migration.** (**A-D**) Disruption of the LINC Complex does not alter migration rate or nuclear transit mechanism in MEFs. (**A**) Representative images of MEFs expressing GFP-Nesprin-2 and either cytoplasmic BFP (BFP Control) or BFP-tagged DN-KASH constructs. Arrows indicate DN-KASH-expressing cells with Nesprin-2 displaced from the nuclear envelope. Scale bar: 20 μm. (**B**) Number of BFP-only or DN-KASH expressing MEFs with Nesprin-2 displaced from the nuclear envelope or non-displaced Nesprin-2. ****, $p < 0.0001$ based on Chi-Squared test between populations. (**C**) Nuclear transit times of MEFs migrating through $\leq 2 \times 5$ $\mu m^2$ constrictions. Red line represents mean, error bars represent standard deviation. Number of cells analyzed per group: $n$ = 35, 68, 25, 77, respectively. (**D**) Nuclear transit mechanism used by MEFs expressing either BFP-only or DN-KASH migrating through $\leq 2 \times 5$ $\mu m^2$ constrictions. Circles represent means from experimental replicates. Number of cells analyzed per group: $n$ = 103 and 102, respectively. Results are based on two independent experiments. (**E-H**) Disruption of the LINC Complex alters nuclear transit rate of non-cortex driven MDA-MB-231 cells. (**E**) Representative images of fixed MDA-MB-231 cells expressing either cytoplasmic BFP or BFP-tagged DN-KASH and immunofluorescently labeled Nesprin-1. Arrows indicate cells with Nesprin-1 displaced from the nuclear envelope. Scale bar: 25 μm. (**F**) Percentage of MDA-MB-231 cells expressing either BFP-only or DN-KASH with endogenous Nesprin-1 displaced from the nuclear envelope. (**G**) Nuclear transit mechanism used by MDA-MB-231 cells migrating through $\leq 2 \times 5$ $\mu m^2$ constrictions. (**H**) Nuclear transit times of MEFs migrating through $\leq 2 \times 5$ $\mu m^2$ constrictions. Red line represents mean, error bars represent standard deviation. *, $p < 0.05$; **, $p < 0.01$; ***, $p < 0.001$, based on one-way ANOVA test. Number of cells analyzed per group: $n$ = 51, 54, 34, 27, 64, 44, 43, 20, respectively. Results collected from three independent experiments.

**Figure S4**

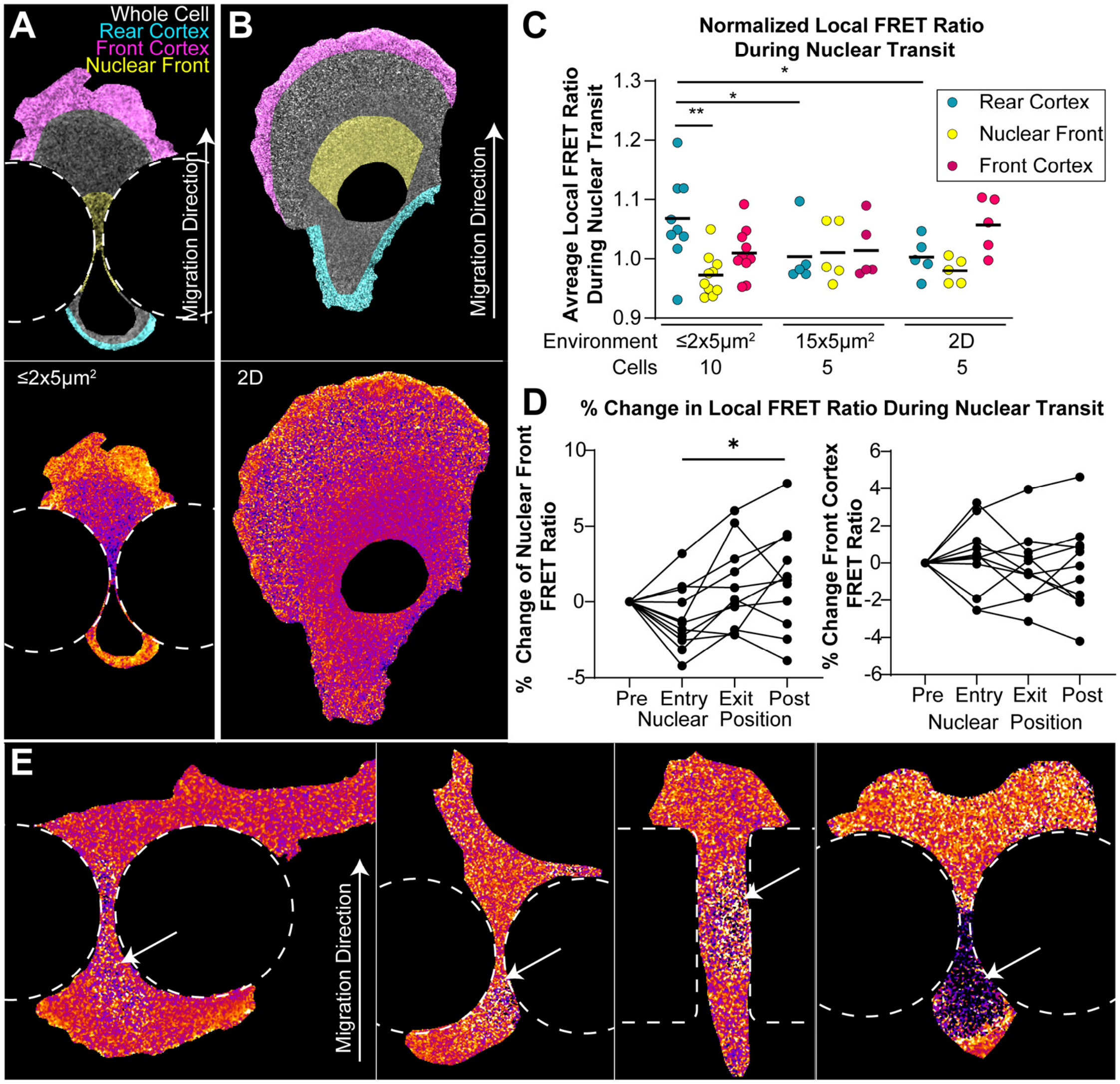


**Supplementary Figure 4. Distribution of active RhoA differs in cells migrating in 2D and 3D**. Representative images depicting localization analysis of RhoA-FRET sensor for cells migrating (**A**) through ≤2×5 μm$^2$ constrictions, or (**B**) on a 2D glass coverslip. Colors in the top image indicate the specific measurement regions defined to represent the "Rear Cortex" (cyan), "Front Cortex" (magenta), and "Nuclear Front" (yellow). Heat maps of measured FRET ratios for each corresponding cell are shown in the bottom row. (**C**) Average FRET ratio in each local region normalized to the average FRET ratio of the whole cell, grouped by environmental conditions. Black lines represent the mean for each group. **, $p < 0.01$ using Friedman test for comparisons within individual cells. *, $p < 0.05$ using unpaired non-parametric Kruskal-Wallis test comparing cells in different conditions. (**D**) Time course of FRET ratio in Nuclear Front and Front Cortex regions in individual cells normalized to the average FRET ratio of the whole cell during each phase of nuclear transit through an individual constriction. (**E**) Representative images of FRET ratios of RhoA-FRET expressing MDA-MB-231 cells without nuclear masking. Arrows point to the location of the cell nucleus.

### Figure S5

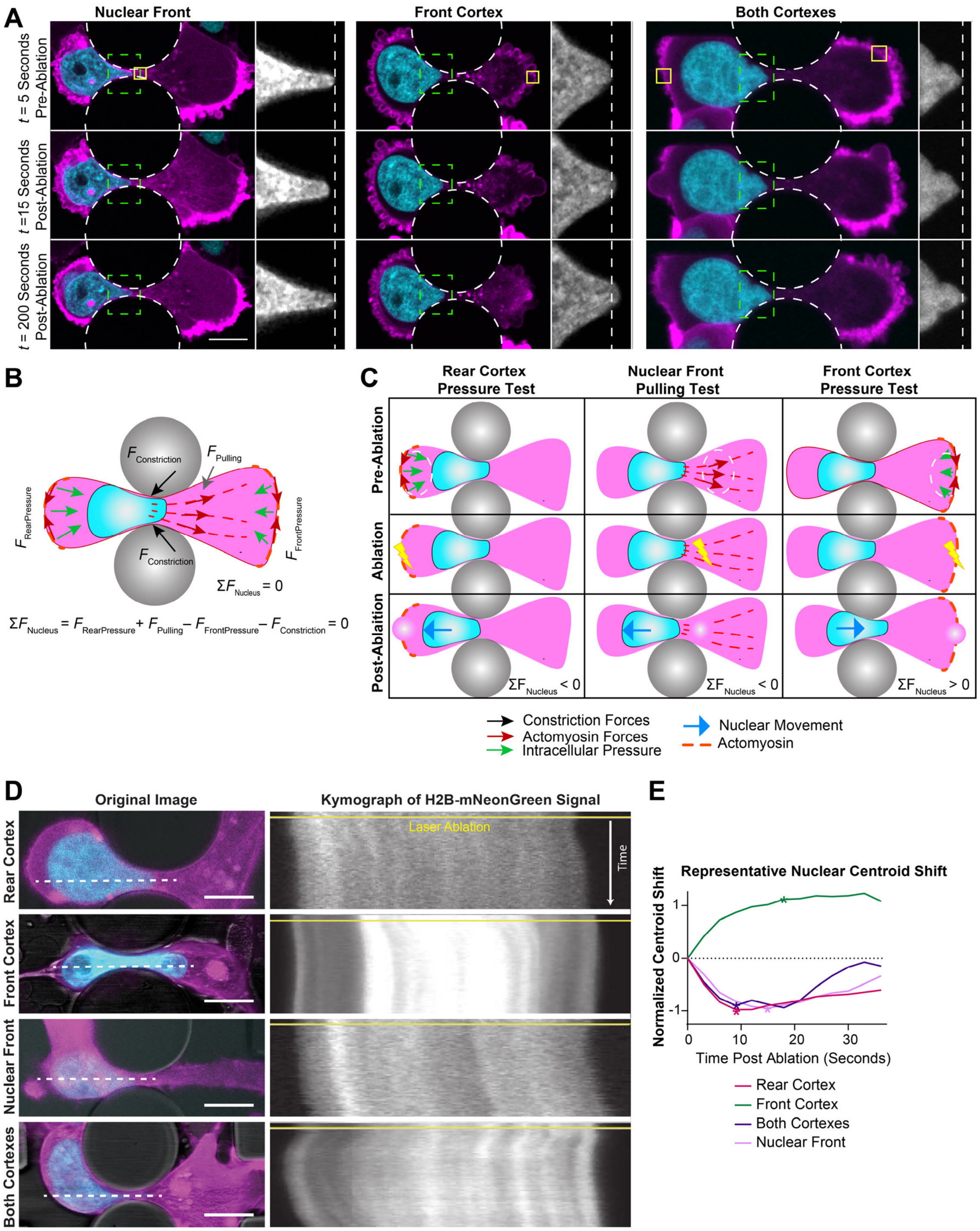


**Supplementary Figure 5. Nuclear movement following laser ablation of specific cytoskeletal regions reveals intracellular forces acting on the nucleus during confined migration.** (**A**) Representative time-lapse sequences of laser ablation experiments for each test group (Nuclear Front, Front Cortex, and Both Cortexes). Yellow box represents region targeted by laser ablation. Dashed white circles represent constrictions. Dashed green box corresponds with inlay of mNeonGreen-H2B Histone signal on right. In inlay region: dashed white line represents starting point of leading edge of nucleus. (**B**) Schematic illustration of forces acting on the nucleus in equilibrium during movement through a constriction. $F_{RearPressure}$ and $F_{Pulling}$ support forward movement of the nucleus, whereas $F_{FrontPressure}$ and $F_{Constriction}$ resist forward movement of the nucleus. While the nucleus is stuck within the constriction, and thus has zero acceleration, the sum of forces acting on the nucleus are zero. If the sum of forces exceeds zero, the nucleus will move forward, if the sum of forces falls below zero, the nucleus will move backward. Green arrows represent forces resulting from intracellular pressure generated by the front and rear cortex. Red arrows represent actomyosin forces. Dotted orange lines represent actomyosin fibers (**C**) Schematic illustration of laser ablation experiment. The laser ablation target regions for each hypothesis to be tested are noted by dashed white circle. The sum of forces acting on the nucleus are assumed to be in equilibrium as the nucleus is stuck in constriction. By eliminating a subset of these forces, the balance of forward and rearward forces is disrupted, and the nucleus will then move in the direction of the removed forces, following the explanation provided for (B). Blue arrow represents predicted movement of the nucleus following ablation of the specified region. (**D**) Nuclear position shifts in response to laser ablation. Kymographs were generated to represent the time course of nuclear position after each type of laser ablation. The dotted line in the images in the left column indicate the trace path for the kymographs. The horizontal yellow line at the top of each kymograph in the right column indicates the time of laser ablation. Scale Bar: 10 μm (**E**) Representative trace of nuclear centroid position over time following laser ablation at each target region. Initial nuclear movement occurs in response to ablation, whereas the subsequent return to the initial position and further advancement shows recovery of actin structures after ablation. Asterisks mark measurement point of peak nuclear shift used for comparison.

### Supplementary Movies

**Movie 1: MDA-MB-231 cells use a cortex-driven mechanism to support nuclear transit through constrictions**. Representative time-lapse sequence of an MDA-MB-231 cell expressing mCherry2-Actin chromobody (magenta) and mNeonGreen-H2B Histone (cyan) transiting through a ≤2×5 $\mu m^2$ constriction by contracting the rear cortex to push the nucleus forward. Scale bar: 10 μm

**Movie 2: MEF cells complete nuclear transit without contracting the rear cortex**. Representative time-lapse sequence of a MEF expressing mCherry2-Actin chromobody (magenta) and GFP-Nesprin-2 (cyan) transiting through a ≤2×5 $\mu m^2$ constriction forming actin fibers that pull at the leading edge of the nucleus. Scale bar: 10 μm

**Movie 3: MDA-MB-231 cells can switch nuclear transit mechanism while passing through constrictions**. Representative time-lapse sequence of an MDA-MB-231 cell expressing GFP-myosin IIA (cyan) and mCherry-myosin IIB (magenta) transiting through a ≤2×5 $\mu m^2$ constriction. In the left panel, the nucleus is outlined in yellow. After initially entering the constriction using a non-cortex driven mechanism (note the low myosin activity in the rear cortex), the cell exits and then re-enters the constriction with elevated myosin activity in the rear cortex, enabling successful nuclear transit. Scale bar: 10 μm
